# Revealing Community Dynamics in Polymicrobial Infections through a Quantitative Framework

**DOI:** 10.1101/2025.09.12.675334

**Authors:** Aanuoluwa E. Adekoya, Carolyn B. Ibberson

## Abstract

Laboratory models provide tractable, reproducible systems that have long served as foundational tools in microbiology. However, the extent to which these models accurately mimic the biological environments they represent remains poorly understood. A quantitative framework was recently introduced to assess how well laboratory models capture microbial physiology in situ. However, applications of this framework have been limited to characterizing the physiology of a single species in human infections, leaving a gap in our understanding of overall microbial community physiology in polymicrobial contexts. Here, we extended this framework to evaluate the accuracy of laboratory model systems in capturing community-level functions in polymicrobial infections. As a proof of concept, we applied the extended framework to a polymicrobial model of human chronic wounds (CW) infection. CWs harbor metabolically diverse bacterial species that engage in a range of microbe-microbe interactions, ultimately impacting community dynamics and disease progression. However, studies on the mechanistic drivers of chronic wound infection have relied on single species or pairwise approaches. Here, we demonstrate that our adapted framework can be used to develop accurate polymicrobial models. Further, we demonstrate that this extended framework can be used to evaluate the occurrence of known microbe-microbe interactions. Building on our prior work in large-scale metagenomic and metatranscriptomic analysis, we propose a highly accurate 6-member synthetic bacterial community model that is representative of the taxonomic and functional complexity of human CW infections. This approach will support the development of ecologically relevant polymicrobial models and the development of better treatment strategies.

## Introduction

Microbiology has long explored the physiology and metabolism of microbes across diverse environments, including within the human body, where direct access is often limited. This progress has been largely enabled by laboratory models that offer tractable systems that can be reproduced and manipulated for in-depth investigation (1–3). While experimental models have proven invaluable for understanding microbial infection physiology, how accurately they mimic the native biological environment they represent is poorly understood. Recognizing the importance of rational laboratory model selection, a quantitative accuracy score framework using RNA sequencing data was proposed and used to improve experimental model systems of cystic fibrosis lung infection and oral infection (1, 4, 5). These studies emphasized the need to investigate and develop laboratory infection models that accurately capture in situ microbial physiology. However, despite the significance of this framework, its application has been limited to characterizing the physiology of a single species in human infection, leaving a gap in our understanding of microbial physiology and overall community dynamics in polymicrobial contexts. Here, we adapt this quantitative framework to evaluate the accuracy of microbial community function in complex polymicrobial infections. We used chronic wound infection as a test case for this approach as a relatively simple, highly relevant polymicrobial infection environment. Further, wounds have multiple features that make them an ideal system to evaluate this approach including accessibility for in situ sampling, previously defined interactions between microbial community members, and measurable outcomes of infection. Chronic wounds (CW) are wounds that fail to heal over a prolonged period (typically beyond six weeks) (6–9). These wounds are characterized by damaged tissues that release extracellular matrix proteins that serve as nutrients and adhesion sites for bacterial colonization (7, 10), and are often hypoxic which allowing for growth of microorganisms with a range of metabolic capacities (7, 11). These metabolically diverse communities establish a range of interactions, which has been shown to impact their survival and overall disease progression (12–17). However, research on microbial drivers of chronic wound infection has primarily focused on single species or pairwise interactions that do not capture the complexity of microbial communities in wound environments. This knowledge gap reduces the extent to which we can compare laboratory findings to patient outcomes in clinical settings.

To address this limitation, we quantified the accuracy of microbial community function in experimental CW models compared to human CW infection and evaluated how modifications to community composition and the host environment impacted experimental model accuracy. We used our community accuracy score framework to assess if known microbe-microbe interactions observed in experimental models are occurring in the infection environment. Further, we leveraged these findings to describe a highly accurate 6-member synthetic wound model system that better reflects the taxonomic and functional complexity of bacterial infection in human CWs. The use of accurate model systems will allow for improved mechanistic studies that capture microbial dynamics in human CW and allow for better correlations of laboratory findings and clinical outcomes.

### Methodology

#### Bacterial Strains for Model and Growth Conditions

Bacterial strains used in this study are listed in Table S1. For the four-member polymicrobial model, these 4 bacteria species were used as previously described (18): 1) *Staphylococcus aureus* (CA-MRSA), 2) *Pseudomonas aeruginosa*, 3) *Finegoldia magna* and 4) *E. faecalis*. For the seven-member community, the following bacteria were incorporated: *Staphylococcus aureus*, *Pseudomonas aeruginosa*, *Streptococcus agalactiae*, *Cutibaterium acnes*, *Finegoldia magna*, *Anaerococcus hydrogenalis*, and *Corynebacterium amycolatum*. *S. aureus*, *P. aeruginosa* and *E. faecalis* overnights were grown at 37°C in Brain Heart Infusion broth (BHI) with shaking at 150RPM (Innova 44 Incubator Shaker series). *C. amycolatum* was grown in BHI + 1% Tween in the same condition. *S. agalactiae* was grown statically at 37°C in Todd Hewitt Broth (THB). *F. magna*, *A. hydrogenalis* and *C. acnes* were grown at 37°C in MTGE anaerobic enrichment broth under anaerobic conditions (Anaerobes Systems AS-150 with 5% Hydrogen and Nitrogen balance) with *C. acnes* being grown for 4 days. For the default four-member and seven-member polymicrobial model, equal volume of each specie at an OD_600_ of 0.1 was used to make 1 ml of the inoculum to have approximately 1 × 10^6^ colony forming units (CFU)/ml. For the 7-member altered inoculum, *P. aeruginosa* was normalized to an OD_600_ of 0.01, *S. aureus* and *S. agalactiae* were normalized to an OD_600_ of OD 0.02 then *F. magna*, *C. acnes*, *C. amycolatum* and *A. hydrogenalis* were normalized to an OD_600_ of 0.1. Thereafter, equal volume of each species was used to make 1ml of the inoculum. Then for each condition, 60ul of the inoculum was inoculated into 540ul of Lubbock Wound-Like Media (19) in 96 well deep well plates (Eppendorf 96/2000 uL) and the plates were grown statically at 37°C in ambient air for 48 hours. At 48 hours, samples were collected and stored in Zymo DNA/RNA shield (Zymo Research R1100-250) for preservation for DNA and RNA extraction. Two technical replicates were combined for each biological replicate.

#### In-vivo model of chronic wound infection

8- to 10-weeks-old female C57BL/6J mice (Jackson’s Lab) were used to perform the murine surgical wound infections as previously described (20–23). Briefly, the OD_600_ of the initially described species were reduced to 0.05 to obtain approximately 5 × 10^5^ CFU in the inoculum for each condition (n = 4). Mice were anesthetized using isoflurane, their backs were shaved and administered bupivacaine. A full thickness 1.5 × 1.5 cm surgical excisional wounds were generated on the dorsal skin, covered with a transparent semi permeable polyurethan dressing. The inoculum for each condition was injected directly under the dressing. At 4 days post infection (4dpi), mice were euthanized, and the wound tissues were excised and immediately saved in 750ul of Zymo DNA/RNA shield (Zymo Research R1100-250) for preservation for DNA and RNA extraction. The protocol for this study was approved by Institutional Animal Care and Use Committee of the University of Tennessee Knoxville (Protocol Number 3071), and all animals were cared for and handled according to the recommendations provided.

#### RNA-sequencing

In vitro and murine wound tissues were collected in DNA/RNA shield as described above. RNA extraction was performed using ZymoBIOMICS DNA/RNA Miniprep Kit (R2002) and the Qiagen RNeasy Mini Kit with Qiazol. From our preliminary experiments, we observed in head-to-head comparisons with the same samples the RNA extracted from these two RNA extraction kits did not significantly impact the resulting sequencing data, as evaluated by clustering and differential expression analyses. The extracted RNA was prepared for sequencing using an Illumina Stranded Total RNA Prep Ligation Kit with Ribo-Zero Plus Microbiome (#20072063) with Illumina Unique Dual Indexes. Sequencing was performed on the Illumina NextSeq2000 platform using a 300-cycle flow cell kit to produce 2×150bp paired reads. 1-2% PhiX control was spiked into the run to support optimal base calling. At least 3 biological replicates of each condition in were used.

#### Quantitative PCR (qPCR) analysis to determine bacterial burden

qPCR analysis was performed to determine bacterial burden at time of sample collection. Briefly, genomic DNA was extracted from invitro and murine samples using the ZymoBIOMICS DNA/RNA Miniprep Kit (R2002) and quantified using Qubit RNA broad range assay (ThermoFisher Q10210) and RNA IQ assay (ThermoFischer Q33221) kits on the Qubit fluorometer (ThermoFisher Q33238). Genomic DNA was also extracted from overnight standards for standard curve quantification. The DNA for each standard curve was diluted 1:10 fold for five dilutions. The universal 16S primers and species-specific primers (Table S2) were used for all bacteria expect for *A. hydrogenalis.* As some of these conditions had numerous bacterial species, end-point PCR assays were performed to test the specificity and cross-reactivity of each primer. Power SYBR (Applied Biosystems Power SYBR™ Green PCR Master Mix) in 20 μl reactions were used for qPCR on Applied Biosystems QuantStudio™ 3 Real-Time PCR System (ThermoFisher) with the following steps 95°C for 10 minutes, and 40 cycles of 95°C for 15 seconds and 60°C for 60 seconds. Unknown bacterial quantification was done in comparison to the standard curve.

#### Human Data Collection

We collected 72 metatranscriptomic datasets identified as being from chronic wounds from the Sequence Reads Archive (SRA) (Table S3). These sequencing files were originally submitted with the project IDs: PRJNA563930, PRJNA726011, PRJNA720438, and SRP135669. Only SRP135669 was not a diabetic ulcer project. To enrich our community composition data, we further analyzed 95 metagenomic datasets. The metagenomic datasets were from PRJNA506988, which had 195 samples (100 of the 195 samples were non-duplicates) and PRJNA610303 (36 samples); however, only 94 were individual samples from chronic wounds. Since the PRJNA506988 study was a longitudinal study, chronicity was determined by a third visit to the facility, which was after 26 weeks of follow-up. All of these were diabetic foot ulcers/infections. All metagenomic samples used are listed in Table S4.

#### Quality Control of Sequencing Data

Using only the forward reads for metatranscriptomic data and paired reads for the metagenomics data, FastQC v0.11.5 (24) was used for quality check, and CutAdapt (25) was used for adapter sequence removal and trimming. SPAdes genome assembler v4.0.0 (26) was used for genome assembly for the metagenome data. Cutadapt v5.0 (25) with universal Illumina adapter AGATCGGAAGAGCACACGTCTGAACTCCAGTCAC was used to remove the adapter sequences and reads with less than 22 bases in length. For the human chronic wounds sample, the trimmed reads (cutadapt output) were mapped to the human genome reference (GRCh38/hg38) using bowtie v2.5.4 (27). The prokaryotic reads were run through SortMeRNA v4.3.7 (28) was used to remove the ribosomal rRNA. (Fig. S1A)

#### Community Composition

MetaPhlAn v4.1 (29), using the Oct2024 SBG CHOCOPhlAn database (CHOCOPhlAnSGB), was used to bin all the prokaryotic reads from the human CW and the in vitro samples to microbes or obtain the community composition data. (Fig. S1A).

#### Community Function Analysis

HUMANn v4.1.1 alpha release (30) with taxonomic profiling from MetaPhlAn v4.1 (29) and other default parameters was used to obtain the metabolic potential data of all the samples in our dataset. The output of these was in UniRef90 IDs and was already normalized for sequencing depth differences to Copies Per Million reads (CPM). We further regrouped the UniRefIDs to EggNOG categories (COG (Clusters of Orthologous Genes) IDs) using the humann_regroup_table function in HUMANn v4.1.1 alpha. The functional categories of these COG IDs were obtained from the eggnog5.0 database FTP (http://eggnog5.embl.de/download/eggnog_5.0/), the COG API linkage on NCBI https://www.ncbi.nlm.nih.gov/research/cog/api/cogdef/, and the KOG database on NCBI https://ftp.ncbi.nlm.nih.gov/pub/COG/KOG/kog. These were used to further annotate the COG IDs obtained in R Studio (R v4.4.1). (Fig. S1A)

#### Accuracy score analysis

To reduce noise in the dataset, as these were short-read RNA Sequencing samples, COG IDs whose cumulative fraction did not contribute to up to 99% of the entire COG ID expression were removed. The quantitative framework (1, 4, 5) was used to obtain the accuracy score of bacteria community functions in these human chronic wound samples for a proper comparison to the model accuracy, and these were referred to as the target samples. That is, to have a baseline for accurate analysis, 2 random human CW samples were selected. The mean expression of each COG ID in these 2 samples and their deviation from the rest of the human CW samples were obtained as a Z score. This step was repeated 1200 times for several possible combinations of samples. The percentage of the COG IDs that fall within 2 standard deviations from the rest of the human CW samples was recorded as the accuracy score (AS_2_) of each sample pair. The average AS_2_ score was calculated based on the number of iterations and was used as the base AS_2_ for comparison for the model to be created. To compare the performance of the in vitro model system to the human CW, the deviation of the mean expression of each COG ID in the in vitro model system from the human CW (target) was obtained. The AS_2_ was the percentage of the COG IDs in the in-vitro model system that fell within 2 standard deviations from the human CW community (the target). This AS_2_ score was then compared to the baseline community AS_2_ score. Additionally, the performance of each major functional category in the model was compared to the baseline community. All statistical analyses were done using RStudio (R v4.4.1).

## Results

### Adapting the Quantitative Framework to a Polymicrobial Context

We first aimed to assess if a previously described accuracy score framework (1, 4, 5) could be applied to evaluate microbial physiology in polymicrobial infection. This approach uses gene expression data to compare bacterial physiology in an experimental model to expression in the actual environment of interest. However, it has been limited to individual bacterial species in human infection (1, 4, 5). Therefore, we adapted this framework to capture overall microbial community function in human chronic wounds (CW). In this framework, gene expression in the environment of interest, here human CW infection, is used as the benchmark. We pulled 72 CW metatranscriptomic samples (Table S3) from three previously published studies (31–33) to serve as the benchmark dataset. We used HUMANn4 (30) to assign prokaryotic reads to UniRef90 gene families as in our previous work (34). The expression of UniRef90 IDs ranged from 68 copies per million (CPM) to 88879CPM with a mean of 14408CPM (Dataset S1A) across the 72 CW samples. Due to the high functional and taxonomic specificity of UniRef90 IDs (35), we aimed to collapse this data to broader functional categories such that gene families with similar functions are grouped together. However, there are several functional grouping schemes, each with varying levels of tradeoffs. Therefore, we evaluated Gene Ontology (GO terms – mean 7121, range 296 – 9745), Cluster Orthologous Genes (EggNOG/COG – mean 3924, range 52 – 11933), KEGG Orthology (KO terms mean 1607, range 14 – 3532), Level 4 Enzyme Classification (Level4EC mean 890, range 16 – 1550) and Protein Family (PFAM – mean 2141, range 42 – 4461) to compare these annotation approaches (Fig. 1A). We found the EggNOG/COG classification provided the best coverage with reduced chance of duplication compared to the other approaches (Fig. 1A & Dataset S1B). We proceeded with the rest of our analysis using the COG classified functional data.

**Figure 1.**
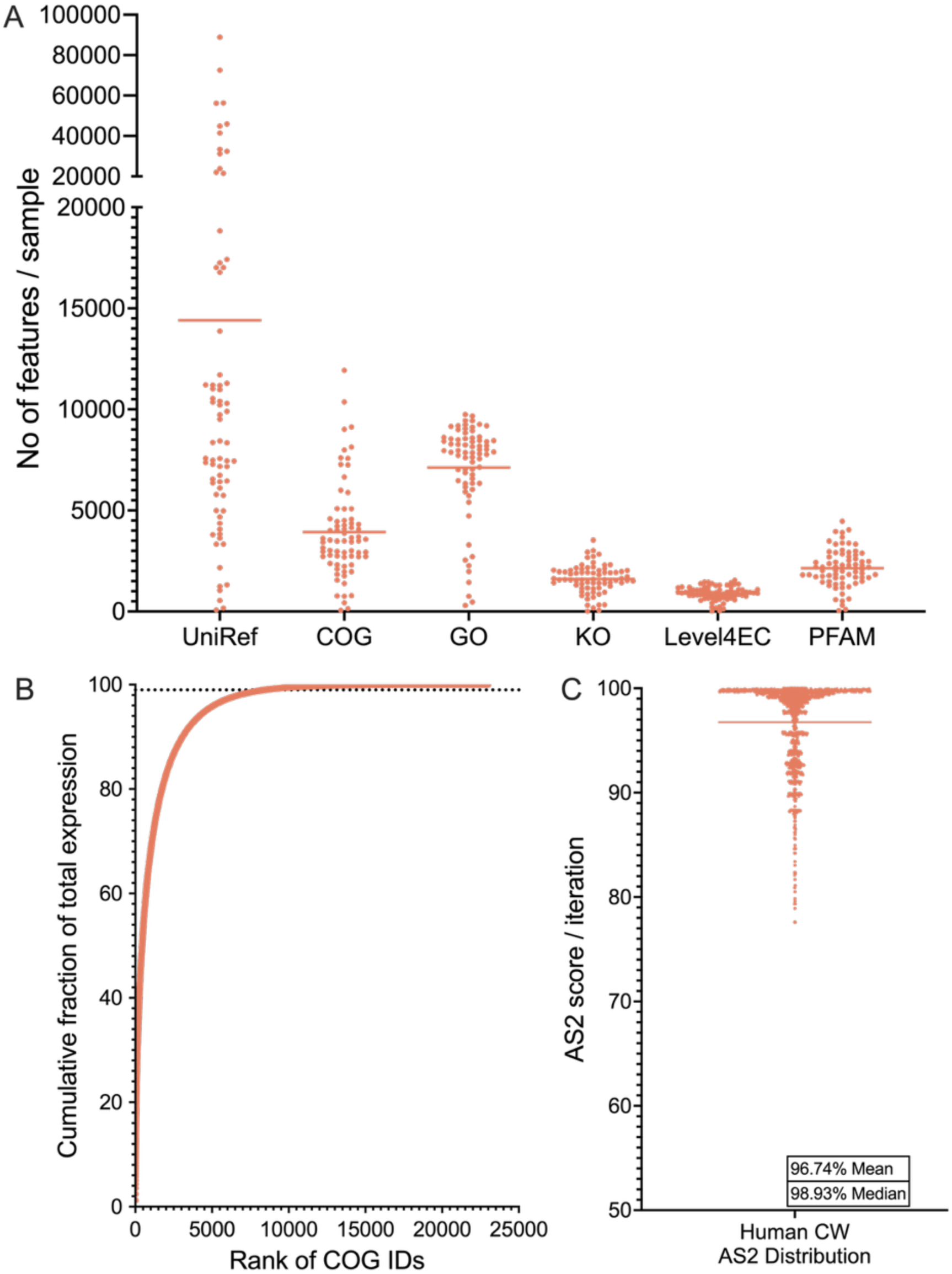
Distribution of functions and accuracy scores in human chronic wound metatranscriptomes. **(A)** Distribution of the different functional categories in human cw samples. Lines at means (UniRef 14408, COG 3924, GO 7121, KEGG 1607, LevelEC 890, PFAM 2141). Each point represents the number of features identified in each human chronic wound sample for each of the annotation schemes. **(B)** Contribution of each COG ID to the total expression across the 66 human CW samples. COG IDs are ordered by their cumulative expression levels. Line at 8486 indicates the point where 99% of the total expression is captured. **(C)** Distribution of the accuracy scores of the subsampled human chronic wound samples across 1200 iterations, Line at mean (96.74%).

Using the COG/EggNOG ID classification we observed an average of 3924 COG IDs per sample, however 6 samples had poor coverage, defined as expression of fewer than 1000 COG IDs and fewer than 2000 UniRef90 IDs (Fig. S1B & Dataset S1A). We removed those samples from subsequent analysis, and the remaining 66 human CW samples were analyzed to identify rarely expressed COG IDs across our dataset. We ranked each COG ID by their expression and prevalence, which revealed that while there were 22986 distinct COG IDs across the 66 samples (Fig. 1B & Dataset S1C), 8486 COG IDs accounted for over 99% of the cumulative expression across all samples, indicating the presence of numerous rarely expressed COG IDs (Fig. S1C & Dataset S1C). Further, these rarely expressed functions accounted for less than 5.5% of the expressed functions in any given human CW sample (Fig. S1D).

Next, we used these 8486 COG IDs as input to calculate the accuracy score (AS_2_) of the human CW samples (Table S3) using a 1200 iteration leave-two-out cross validation approach. We found the AS_2_ were between 77.59% and 100%, with a mean of 96.74% and a median of 98.93% (Fig. 1C). This demonstrates the 66 human CW samples have high similarity, irrespective of the patient, sample collection method, processing, and RNA sequencing protocol. This mean is above the 95% confidence interval and sets the upper limit or benchmark for comparison.

### The Quantitative Framework can be Used to Assess Microbial Dynamics in a Polymicrobial Model

To validate this adapted framework, we employed a previously published 4-member polymicrobial laboratory model of human CW infection (18). We seeded *S. aureus*, *P. aeruginosa*, *E. faecalis* and *F. magna* into wound-like media (WLM) (19), a well validated media for in vitro studies of human CW infection and obtained transcriptomic data at 48 hours post inoculation. We used the annotation approach highlighted above to identify community functions and calculate the accuracy score of the expression of these COG IDs when compared to the human CW. We found the average number of COG IDs per sample was 3860, like the human CW samples (3461) (Fig. 2A), however the overall AS_2_ was 77.98% (Fig. 2B). Looking at the broader functional categories, we saw that COG IDs related to “virulence” were the least accurate (AS_2_ 64.10%, Fig. S2) in the 4-member polymicrobial model. Further, “metabolism” and “cellular processes and signaling” were also not well captured with AS_2_ of 86.65% and 89.87%, respectively (Fig. 2B). While the overall accuracy score of the 4-member polymicrobial model is significantly lower than the benchmark, this data demonstrates the quantitative framework can be adapted to polymicrobial contexts and reveals functional categories that can be targeted to improve the accuracy of the model system.

**Figure 2.**
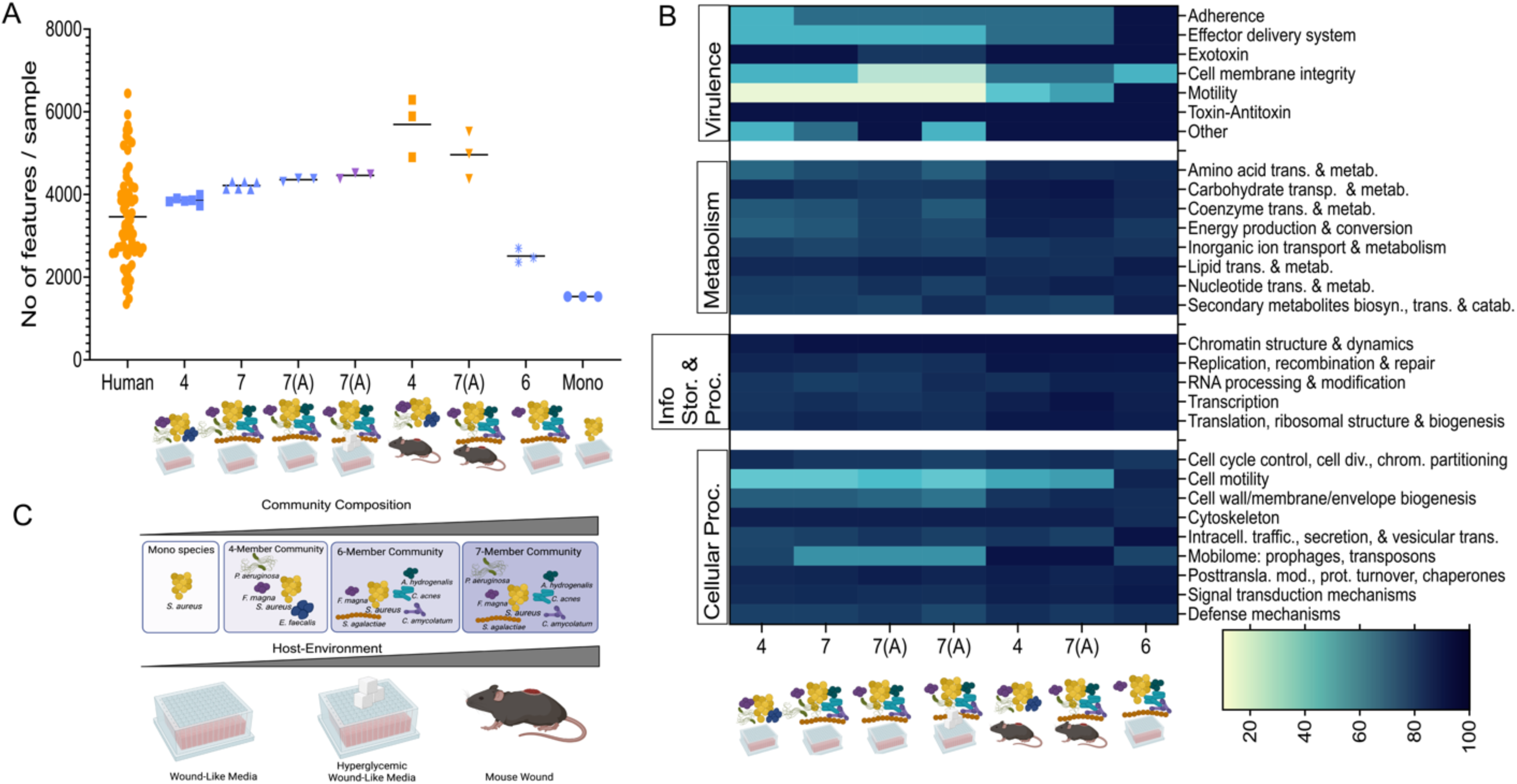
Community composition and host environment are key drivers of accuracy scores. **(A)** Distribution of COG IDs in all samples evaluated for accuracy score. Line at mean (Human 3461, 4-member WLM 3860, 7-member WLM 4218, 7-member altered WLM 4359, 7-member altered hyperglycemic WLM 4462, 4-member murine 5694, 7-member murine 4961, 6-member WLM 2511, *S. aureus* monoculture WLM 1531). **(B)** Heatmap of the accuracy scores of each subcategory. in the samples evaluated with the overall accuracy scores being 4-member WLM 77.98%, 7-member WLM 78.00%, 7-member altered WLM 82.02%, 7-member altered hyperglycemic WLM 77.62%, 4-member murine 89.70%, 7-member murine 90.78%,, 6-member WLM 96.27%, *S. aureus* monoculture 89.80%. **(C)** Two axes that can be manipulated iteratively to improve the overall accuracy of our experimental model system. Abbreviations: 4 = 4-member, 7 = 7-member, 7(A) = 7-member with altered inoculum, 6 = 6-member, Mono = *S. aureus* monoculture in any condition.

### A Seven Member Community Captures Taxonomic Diversity in Human CW

Next, we considered approaches to rationally improve the accuracy of this experimental model and proposed two key axes we could manipulate: community composition and host environment (Fig. 2C). We first aimed to modify the community composition therefore sought out to understand the taxonomic composition of human chronic wounds and build a representative synthetic community. We reanalyzed the microbial community composition of the 72 human CW meta-transcriptomes with MetaPhlAn4 (36) to determine the taxonomic diversity. Of these 72 samples, MetaPhlAn4 could determine community composition in 58 and a total of 183 species were identified. However, only 141 species had a relative abundance greater than 0.1% in a least 1 sample. We then calculated the cumulative abundance for each species, that is considering the abundance for all species across all samples as 100%, how much did each species contribute. We found only 46 bacterial species made up over 95% of the cumulative abundance across all samples (Fig. 3A, Dataset S2A), and only 19 bacterial species comprised over 80% of the data (Fig. 3B, Dataset S2A). These highly abundant and prevalent species were a mix of canonical pathogens such as *S. aureus*, *S. agalactiae*, *Staphylococcus epidermidis*, *Escherichia coli*, and *P. aeruginosa,* and commensal anaerobes such as *Cutibaterium acnes*, *Anaerococcus obesiensis*, and *F*. *magna*. Further analysis revealed that *S. aureus* was the most abundant with *F. magna* the most prevalent (Fig. 3A&B).

**Figure 3:**
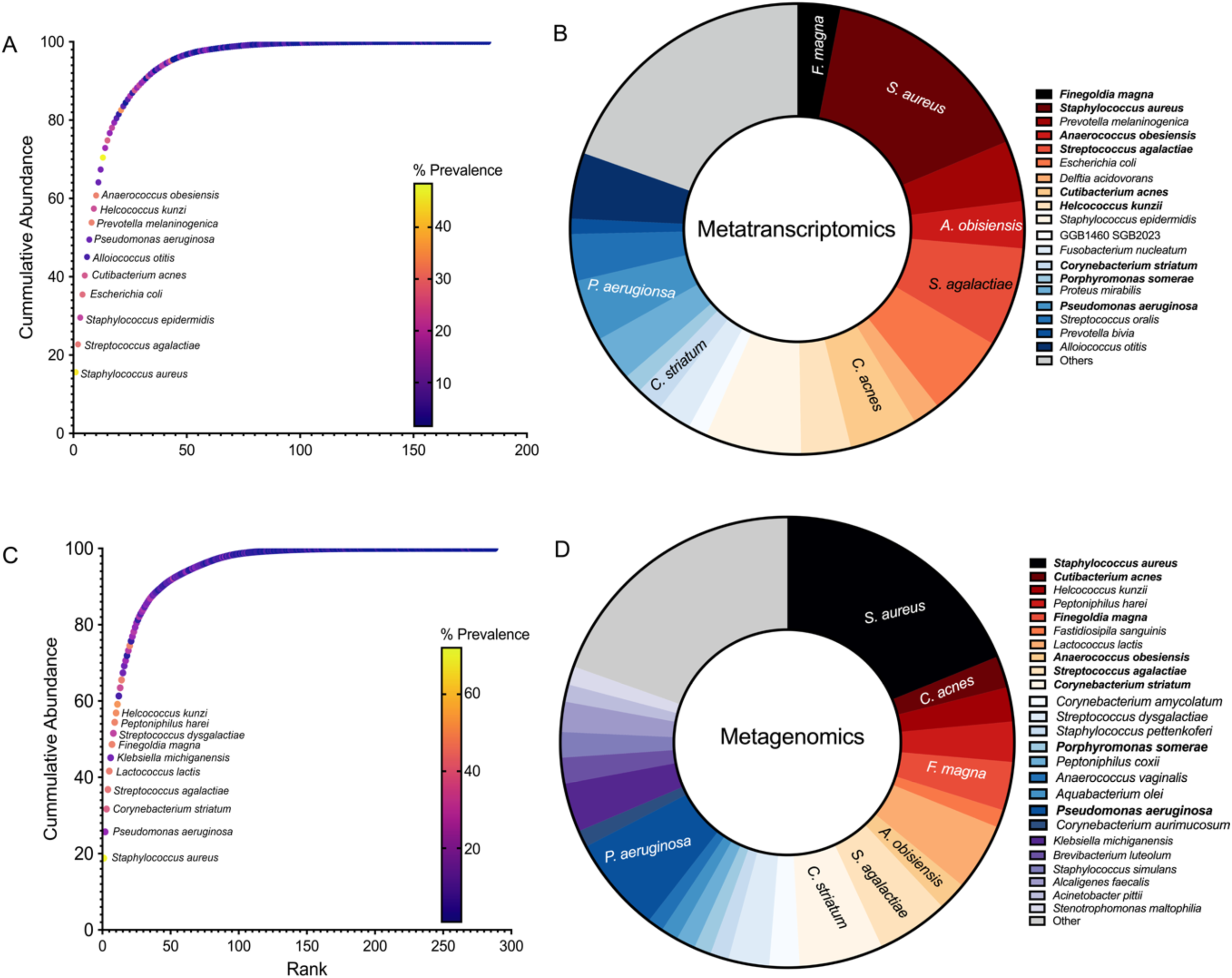
Only a few species comprise the key species in the metatranscriptomic and metagenomic datasets. Graphs show the cumulative abundance of bacterial species in the publicly available data that were evaluated. **A&B** Metatranscriptomic data showing **(A)** all the species ranked by abundance and colored by prevalence with approximately only top 60% labeled and **(B)** all species colored from warm to cool by prevalence and size of wedge is abundance highlighting top 80% with distinct colors and other species grouped as “other” colored in grey. **C&D** Metagenomic data showing **(C)** all the species ranked by abundance and colored by prevalence with approximately only top 60% labeled and **(D)** all species colored from warm to cool by prevalence and size of wedge is abundance highlighting top 80% with distinct colors and other species grouped as “other” colored in grey.

To further understand CW community composition, we identified an additional 94 human CW metagenomic samples (Table S4) from previously published studies (37, 38). Using the approach above, we identified 288 total species, however similar to the metatranscriptomic data, 68 species comprised 95% of the total abundance (Fig. 3C, Dataset S2B) again demonstrating the presence of many bacterial taxa with low abundance and prevalence. Further, we found 25 species contributed to over 80% of the cumulative abundance (Fig. 3D, Dataset S2B). These top species were also a mix of canonical pathogens, such as *S. aureus*, *S. agalactiae*, *E. coli*, *P. aeruginosa,* and commensals such as *Corynebacterium amycolatum, C. acnes*, *A. obesiensis*, and *F*. *magna*. Further analysis revealed that *S. aureus* was the most abundant and the most prevalent (Fig. 3C&D). Interestingly, there was a high degree of overlap between the metagenomic and metatranscriptomic datasets, with 9 species including *S. aureus, C. acnes, Helcococcus kunzii, F. magna, A. obesiensis, S. agalactiae, Corynebacterium striatum, Porphyromonas somerae, and P. aeruginosa* (Fig. 3B&D, Dataset S2C), to be highly abundant and prevalent in both datasets despite no sample or clinical overlaps.

Building off this data, we seeded a 7-member mock community in WLM consisting of *S. aureus, C. acnes, F. magna, A. hydrogenalis, S. agalactiae, C. amycolatum, and P. aeruginosa,* as publicly available representatives of genera which were highly abundant and prevalent in both datasets and obtained transcriptomic data at 48 hours post inoculation. We targeted a seven-member community as multiple studies have found chronic wounds to contain ∼6-7 species per wound (11, 34, 39, 40) with diverse methodology. We found the number of expressed COG IDs in the 7-member community was higher than the 4-member mock community (Fig. 2A). However, the 7-member community had an AS_2_ of 78.00% (Fig. 2B & Fig. S2), demonstrating that while our 7-member community better captured the taxonomic diversity and number of expressed COG IDs compared to the human CW, it did not accurately replicate microbial community function. Specifically COG IDs in the categories “virulence”, “metabolism”, and “cellular processes” were poorly captured with AS_2_ scores of 69.23%, 87.7% and 89.97%, respectively (Fig. S2).

### Microbe-Microbe Interactions Impact the Stability of Community Members and Overall Community Function

Microorganisms in infection environments exhibit several interactions that impact disease progression and overall community physiology (12, 15, 20, 21, 41–43, 43–46). Recognizing that interactions between community members may be impacting the accuracy score, we highlighted two well studied members, *S. aureus* and *P. aeruginosa*, for further investigation. Several studies have shown *P. aeruginosa* products such as LasA protease, 4-hydroxy-2-heptylquinoline-N-oxide (HQNO) and phenazines support the killing of *S. aureus* in vitro (47–50). Therefore, we hypothesized we could use the accuracy score framework to evaluate if these known interactions were occurring in situ. We predicted if this antagonistic interaction was occurring in human CW, the accuracy score of the model should not decrease when *P. aeruginosa*-*S. aureus* competitive interactions were known to occur in vitro (48, 51). To test this prediction, we first modified the inoculation strategy of the 7-member community in WLM to increase the ratio of *S. aureus* to *P. aeruginosa* and give *S. aureus* a competitive advantage in the inoculum. Importantly, qPCR showed all members of the community were present up to 48 Hours with this inoculation strategy (Fig. S3A). Interestingly, we found this modified inoculum community had an AS_2_ of 82.02% (Fig. 2B & Fig. S2), indicating that lowering the ratio of *P. aeruginosa* compared to other community members increased the accuracy of the experimental model. More importantly, COG IDs supporting “metabolism” and “cellular processes” were increased, with “virulence” and “information storage and processing” slightly reduced when compared to the initial 7-member community set up (Fig. 2B & Fig. S2). While this modified inoculation strategy increased the accuracy score, a closer look into the viability of *P. aeruginosa* and *S. aureus* with plate counts revealed that *P. aeruginosa* grows rapidly when compared to *S. aureus* and that the burden of *S. aureus* was overall reduced in this model (Fig. S3B), demonstrating that reducing the amount of *P. aeruginosa* in the inoculum was not sufficient to increase the stability of *S. aureus*. Additionally, this data supports the limitation qPCR data is not an absolute quantification of live bacterial cells. Subsequent experiments with the 7-member community were performed with this altered inoculation strategy.

### The Host Environment Strongly Impacts Microbial Physiology

While modifying community composition is one way to improve model accuracy, microbes are known to be highly responsive to their environments (12, 52–55). Therefore, we predicted we could instead improve model accuracy by modifying the growth environment (Fig. 2C). Approximately 97% of the human CW metatranscriptomes that comprise our benchmark were from diabetic foot ulcers. Therefore, we predicted modifying WLM to be hyperglycemic may better reflect the infection environment and improve model accuracy. We also predicted the increased glucose may also increase *S. aureus* fitness in the 7-member community, as several studies have shown S. aureus has multiple glucose transporters that it uses to increase its glucose import from its extracellular environment and that this increases *S. aureus* virulence (53, 56–58). We seeded the altered 7-member mock community in hyperglycemic WLM (4.5 g/L glucose) consistent with diabetic blood glucose levels (59–62). We found the hyperglycemic model had an AS_2_ of 77.62% and that “metabolism”, “virulence”, and “cellular processes” had lower accuracy scores in the hyperglycemic model compared to non-hyperglycemic WLM (Fig. 2B & Fig. S2). Further, we found while the addition of glucose somewhat rescued *S. aureus* up to 24 hours post inoculation, at 48 hours *S. aureus* had fallen to approximately 1X10^4^ CFU/ml while *P. aeruginosa* had grown to 6 × 10^9^ CFU/ml (Fig. S3B). This data is suggestive of *P. aeruginosa* killing *S. aureus* in WLM, similar to previous observations in vitro (48, 49, 51, 63).

To further investigate the impact of the host environment on overall community function we infected a murine surgical chronic wound model of infection (20–23) with the 4-member and altered 7-member mock communities. We found at 4 days post infection the mean expressed COG IDs of these synthetic communities was 5696 and 4961, respectively (Fig. 2A), and the accuracy scores were 89.70% and 90.78%, respectively (Fig. 2B & Fig. S2). Further, we found the “virulence”, “metabolism”, and “cellular processes” categories which were poorly captured in the in vitro experimental models analyzed had improved accuracy scores (Fig. S2), highlighting the critical role of the infection environment in shaping microbial metabolism and physiology, and in turn, driving overall community function.

### Accuracy of Cellular Processes and Virulence across Laboratory Models

The accuracy score analysis revealed microbial community function across different community compositions (4member vs 7member) and environments (WLM, hyperglycemic WLM, and murine wounds) did not accurately mimic in situ microbial physiology. Therefore, we wanted to investigate the role of individual functions on model accuracy. We found the meta-category “virulence” was the least accurate category in the 4-member WLM condition and gradually increased in accuracy across iterations of model improvement (Fig. 2B). Within the “virulence” category, the subcategories “effector delivery system”, “adherence”, “cell membrane integrity” and “motility” were the least accurate. For the “effector delivery system” subcategory, COG IDs involved in the Type II, Type VI (T6SS) and Type VII (T7SS) secretion systems (COG4796, COG3157 and COG4499) had significantly higher expression in the in vitro models compared to the murine and human samples (Fig. 4A). Further, we found expression of these COG IDs in the murine models was more like the human CW samples. Similarly, COG IDs involved in “adherence” and “motility” (pili: COG3166, COG3539; and flagella: COG1334, COG1344, COG1345, COG1419, COG1516, and COG1536) had significantly higher expression in the experimental models, particularly the in vitro models, compared to the human CW (Fig. 4B). Functions involved in “cell membrane integrity” (COG0744, COG0768, COG5009 and COG2885) also displayed significantly higher expression in vitro compared to our human CW samples (Fig. 4C). In contrast, the “mobilome” subcategory within the “cellular processes” category was also inaccurate and displayed higher expression in the human CW samples compared to the laboratory models (Fig. 2B). Particularly, multiple transposases (COG2826, COG3436, and COG3039) had significantly higher expression (p-value < 0.05) in the human samples compared to our experimental models (Fig. 4D). Of note, many inaccurate functions are COG IDs encoded solely by *P. aeruginosa* in our synthetic communities, suggestive that inclusion of *P. aeruginosa* in these communities could be driving the reduced accuracy.

**Figure 4:**
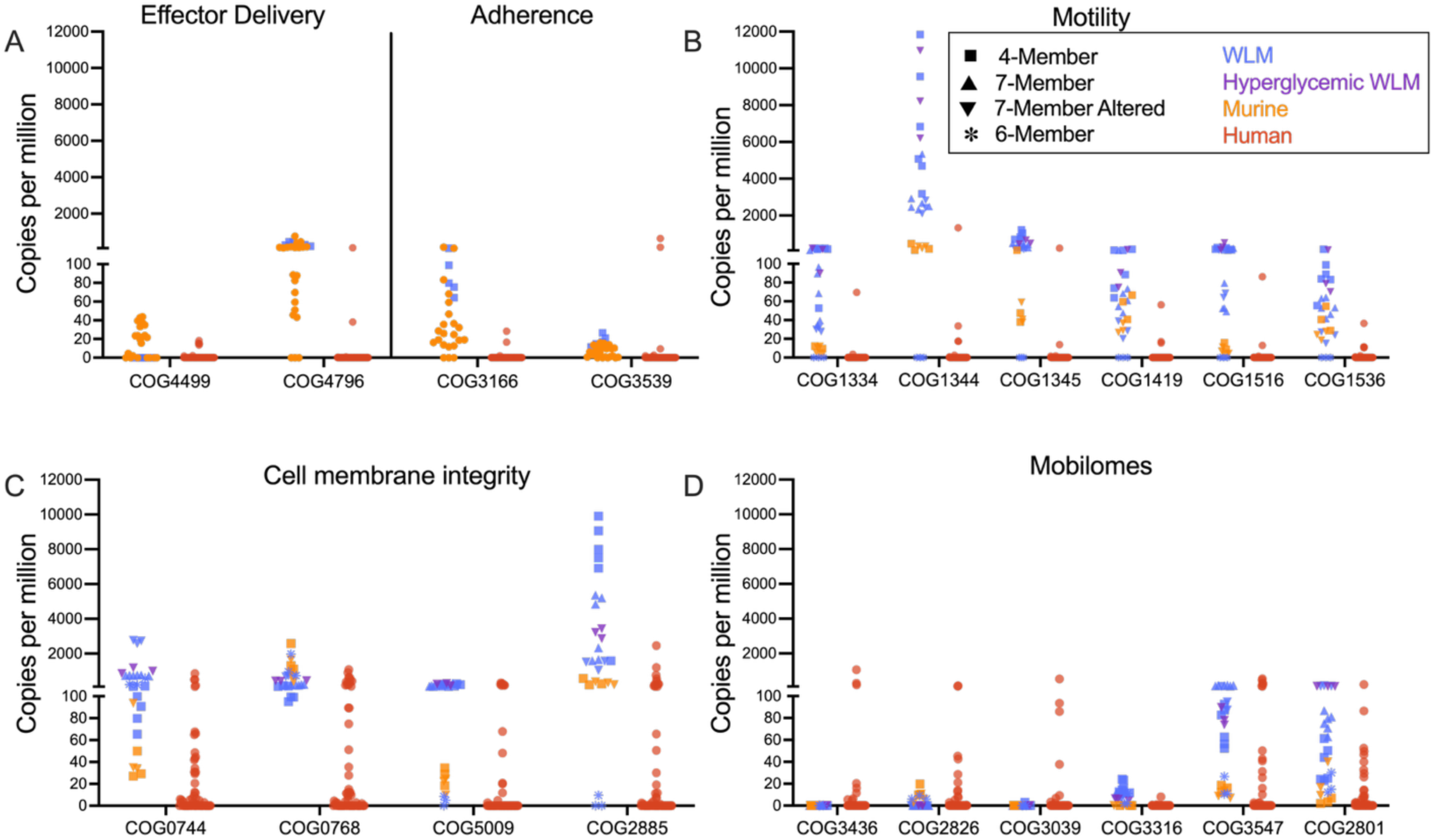
Virulence related functions have higher expression in model samples while genetic mobility is higher in human samples. Graphs show the expression of COG IDs that drive accuracy scores in all the samples evaluated. Y-axis for A-D is the normalized expression (copies per million) for each COG ID. Each point is an individual sample. COD IDs are grouped by subcategories they belong to. Samples are grouped as humans or model. Model samples are shaped by community structure and colored by infection environment.

### A Six-member Polymicrobial Community Better Capture Insitu Microbial Physiology

We observed in our analyses that the presence of *P. aeruginosa* both competitively reduced the burden of *S. aureus* and drove key functions towards reduced accuracy. Therefore, we hypothesized the elimination of *P. aeruginosa* in the 7-member community model would increase model accuracy. We inoculated WLM with a 6-member community comprised of all members of our described 7-member community, but with *P. aeruginosa* omitted, and calculated the accuracy score 48 hours post inoculation as above. While we found there was a significant decrease in the mean number of COG IDs identified in the 6-member community (2511) compared to our previous models (Fig. 2A), the AS_2_ dramatically improved to 96.27% (Fig. 2B & Fig. S2). To verify that this improvement was not driven solely by *S. aureus*, we compared the 6-member community to *S. aureus* alone in WLM at 48 hours, which yielded fewer expressed COG IDs (mean 1531; Fig. 2A) and a lower AS_2_ (89.80%; Fig. S2), indicating that *S. aureus* alone is not sufficient to capture community physiology in human CW infection. Instead, the reduced number of COG IDs in the 6-member community likely reflects the smaller genome sizes of its members.

We next examined how the 6-member community achieved higher accuracy. Principal component analysis (PCA) revealed the 6-member community and murine models clustered closer to some human CW samples than the in vitro models (Fig. S4A&B), and expression of the top 50 COG IDs driving clustering was more similar between the 6-member community and human CW samples than between the in vitro models (Fig. S4C). Additionally, “virulence” and “cellular process” categories, which were poorly captured in earlier models, improved to 94.87% and 96.24%, respectively (Fig. 2B). Notably, the “motility” sub-category in the 6-member community matched human CW expression levels, suggesting that *P. aeruginosa* had previously driven motility and CW communities are largely non-motile. In contrast, functions involved in “cell membrane integrity” remained poorly captured in all experimental models, but particularly in vitro conditions (Fig. 4C). Finally, “mobilome” functions had reduced expression and accuracy (87.5%) relative to both human CW samples and murine models (Fig. 4D).

## Discussion

Laboratory models remain essential tools, particularly in pathogenesis research where direct studies are constrained by ethical limitations. Therefore, it is critical to understand the accuracy of experimental models in capturing microbial physiology the native environment. Recent studies (1, 4, 5) have reported the accuracy of some models, however, none of these have been done in a polymicrobial context. Here, we provide the first quantitative evaluation of the accuracy of polymicrobial infection model systems using human CW, a known polymicrobial human infection (11, 34, 39, 40). With this extended framework, we can evaluate the accuracy of polymicrobial models in any biological system, and more importantly we can use this extended framework to develop more representative polymicrobial models.

We demonstrated that while a well 4-member polymicrobial model of human CW accurately captured some aspects of microbial CW community functions related to “virulence” and “cellular processes” were poorly represented. We used this as a baseline for iterative experimental model improvement along two axes: community composition and host environment. We leveraged publicly available metatranscriptomic (31–33) and metagenomic (37, 38) datasets from human chronic wound to understand the taxonomic diversity of human chronic wounds and applied this knowledge to build a representative synthetic community. Further, we used a range of culture and in vivo conditions to better understand the impact of the host environment on microbial community function.

An interesting trend in our data was that “virulence”, and “cellular processes” were the least accurate categories across all model types, and that these gradually improved across iterations (Fig. 2B). We focused on the least accurate sub-categories including “effector delivery system”, “adhesion”, “motility”, “cell membrane integrity”, and “mobilomes” for further exploration into what was driving this inaccuracy. Within the “effector delivery system” sub-category, COG IDs involved in the Type II (T2SS), Type VI (T6SS) and Type VII (T7SS) secretion systems contributed to this reduced accuracy and are known to be involved in bacterial competition (12, 64–68). The high expression of the T6SS and the T7SS in the in vitro polymicrobial models in this subcategory (Fig. 2B & Fig. 4A) could indicate an over stimulation of bacterial competition in the models compared to the human CW environment. Interestingly, this subcategory had improved accuracy in the murine models and in the 6-member in vitro model, suggesting *P. aeruginosa* is contributing to that sub-category and that expression of these systems is reduced in vivo.

Similarly, the “adhesion” and “motility” sub-categories had significantly higher expression of COG IDs related to the bacterial Type VI Pilus and flagellar systems in the in vitro communities compared to the human wound samples (Fig. 2B and Fig. 4A&B). *P. aeruginosa*, the sole member of our synthetic communities that encodes for these functions, uses its Type IV Pilus and flagellar systems for motility and to sense surfaces for possible attachment. The high expression of these functions in the in vitro models but not the murine samples indicate that while *P. aeruginosa* possess this capacity, it likely does not express these structures at high levels in vivo in wounds. The *Pseudomonas* pilin has been shown to activate immune response (69, 70) and the flagellar system is known to activate the host immune system through the Toll-like Receptor 5 (70–72). The high expression of these systems in vitro but not in vivo could suggest the cells are highly motile in vitro and that *P. aeruginosa* deduces expression of these systems in vivo, possibly to avoid activation of host immunity.

Interestingly, we observed multiple COG IDs involved in “cell membrane integrity” (COG2885, COG0744, COG0768, and COG5009) had significantly higher expression in our experimental models compared to the human CW samples (Fig. 4C). The high expression of these membrane integrity proteins in our in vitro and murine laboratory models could be reflective of the stressful environment the cells are in. Membrane integrity related functions often participate in key processes such stress response or homeostasis for bacterial survival (73, 74). In vitro the cells are in batch culture, allowing for the accumulation of metabolic waste products and pH shifts. Additionally, we found evidence of competitive interactions in these models, further increasing bacterial stress. In addition, the mice have a fully active immune system the microbes need to overcome. This contrasts with human CW infections, which are known to have impaired immune responses (75–81) and our data indicates lower expression of functions associated with microbe-microbe interactions. Therefore, the CW environment may induce lower bacterial stress responses relative to our experimental models.

While multiple inaccurate virulence subcategories were driven by overexpression in the experimental models compared to human CW infection, the sub-category “mobilomes” had significantly higher expression in the human CW samples compared to the laboratory models (Fig. 4D), driven largely for COG IDs for multiple transposases (Fig. 4D). Higher transposase expression in vivo may reflect increased selective pressures in the dynamic wound infection environment which may require increased genomic flexibility to survive against the combined stressors of host immunity, nutrient limitation, and polymicrobial competition.

One interesting application of our polymicrobial accuracy score framework is to evaluate if known microbe-microbe interactions occur in human infection by their impact on model accuracy. Several studies have shown antagonistic interactions between *P. aeruginosa* and *S. aureus* in vitro through the production of *P. aeruginosa* exoproducts (47–50, 63). In support, we found increases in *P. aeruginosa* burdens in our in vitro models were accompanied by decreased *S. aureus*, and the models where we observed this trend had lower accuracy (Fig. 2B & Fig. S3B). In contrast, the murine models where previous work has shown co-existence of *S. aureus* and *P. aeruginosa* (21) had improved accuracy (Fig. 2B). The observation that conditions with reduced *S. aureus* and higher abundance of *P. aeruginosa* were less accurate suggested that a competitive interaction was occurring in those in vitro models and that this interaction is not a key driver of microbial community function in human CW infections. Supportive of this idea, the omission of *P. aeruginosa* in the 6-member community was accompanied by the increased bacterial burdens of *S. aureus* (Fig. S3B) and an increased accuracy score that matched the mean and median of the human CW accuracy score (Fig. 2B). Importantly, the 6-member WLM condition had a better coverage of functions that were poorly captured in the previous models (Fig. 4A-D) further showing that reducing the expression, here by omission, of those functions was key to mimicking in situ microbial physiology. While it is unlikely that the human chronic wounds community lack *P. aeruginosa*, this data suggests that the competitive advantage of *P. aeruginosa* in vitro does not mimic the human CW infection environment. Future studies will seek to better understand the molecular mechanisms driving model accuracy. Together, these will increase our understanding of the microbial community functions in human CW infections and provide insights into the microbe-microbe interactions that occur in situ.

While we believe our polymicrobial accuracy score approach is a powerful tool for understanding the complex biological processes and interactions within polymicrobial communities, there are some key limitations. We selected COG classification in our approach because it provided the best balance with trade-offs among the annotation schemes tested. However, this choice in annotation strategy does have trade-offs. GO and KO terms offer detailed functional information but are biased towards well-annotated organisms, and for GO terms functions are collapsed into only three broad categories with frequent overlaps making the interpretation challenging. Level4EC focuses on enzymes with known catalytic activity, limiting its utility for uncharacterized proteins. PFAM enables domain-based predictions but showed limited coverage as few UniRef90 IDs had direct PFAM IDs conversions. In contrast, COG classification grouped orthologs across taxa to into hierarchical functional for further traceability and extensive analysis. However, a key limitation is COG databases are not always up to date, leaving many IDs poorly characterized. These challenges highlight the need for improved annotation resources as accurate annotation is critical for translating omics data into biological insights. Our 7-member synthetic community included bacterial isolates representative of most abundant and prevalent genera in human CWs. However, due to isolate availability, we substituted *Anaerococcus obesiensis* with a closely related publicly available species, *A. hydrogenalis*, and substituted *Corynebacterium striatum* with a publicly available *C. amycolatum* isolate in our synthetic community. While both species were also found in our datasets, this is also a key limitation.

In conclusion, we demonstrated the application of an extended framework to develop accurate polymicrobial models in any system. Further, we propose a 6-member synthetic bacterial community that captures the taxonomic diversity and functional complexity of human chronic wound infections that can be applied to future work. Importantly, our framework can be used to evaluate if known in vitro microbe-microbe interactions occur in situ. Collectively, this approach is an important next step towards accurate model systems that reflect the biological processes they are meant to capture.

## Data availability

The key codes involved in the analyses done for this manuscript can be found on our GitHub page via this link https://github.com/Ibberson-Lab/Assessing-Microbial-Community-Physiology-in-a-Polymicrobial-Infection. The raw sequencing files for the new RNAseq data collected have been submitted to SRA under accession number PRJNA1314253.

## Acknowledgments

We would like to thank Marvin Whiteley, Alexander Horswill, and Brandon Kim for generously sharing bacterial strains for this work. We would like to thank all members of the Ibberson lab for thoughtful feedback in the development of this manuscript. C.B.I is supported by the National Institutes of Health (R35GM160019).

---

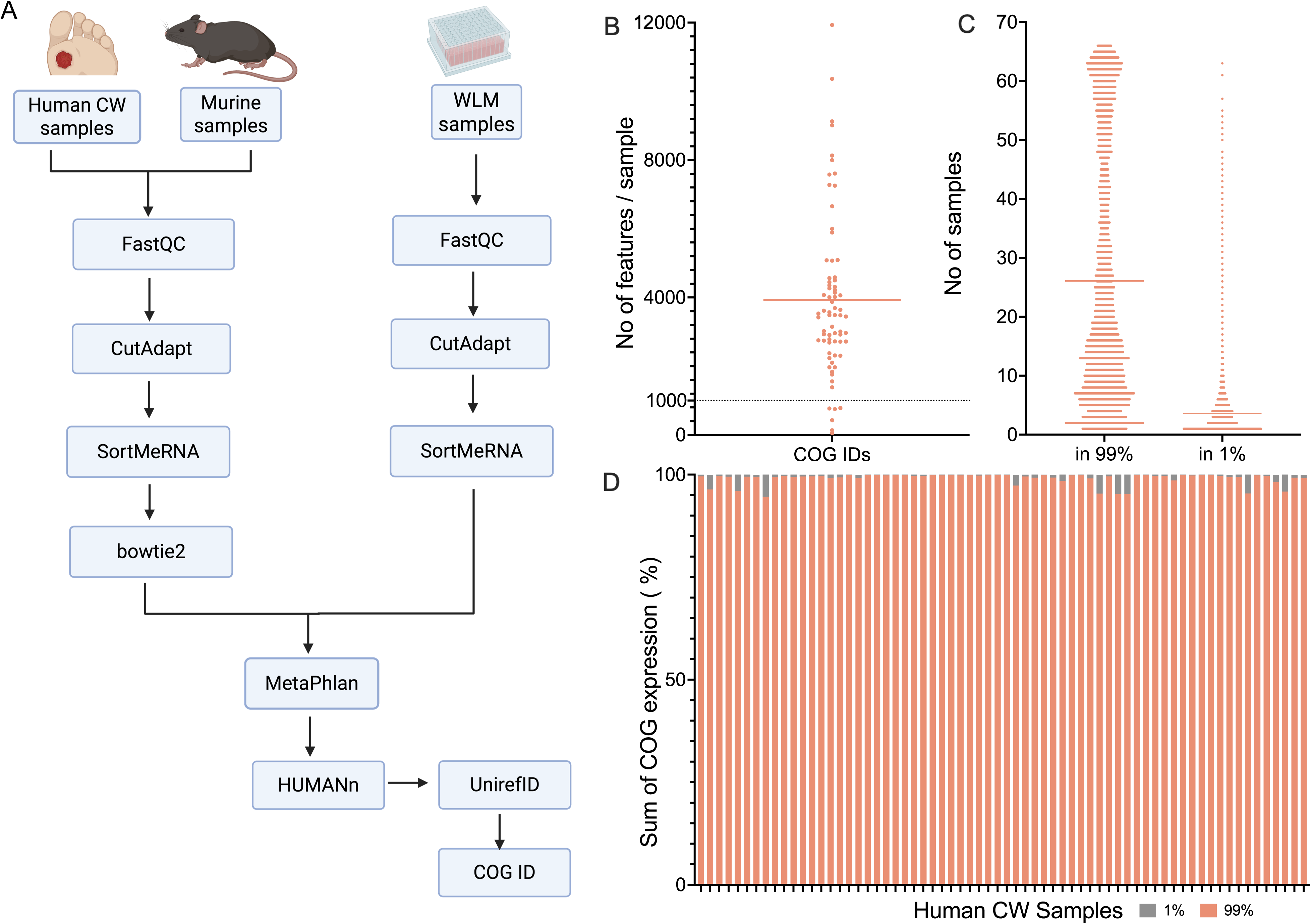

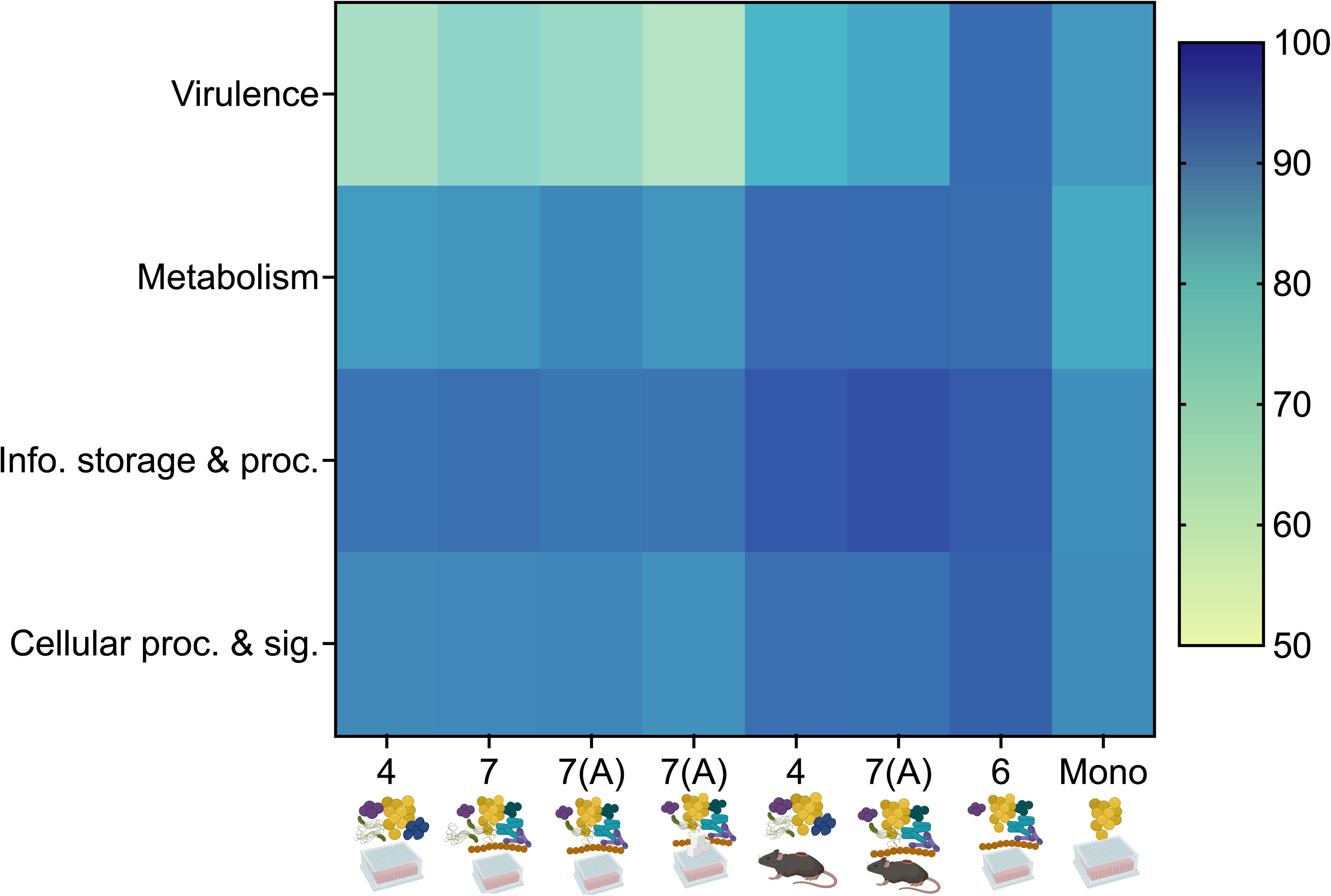

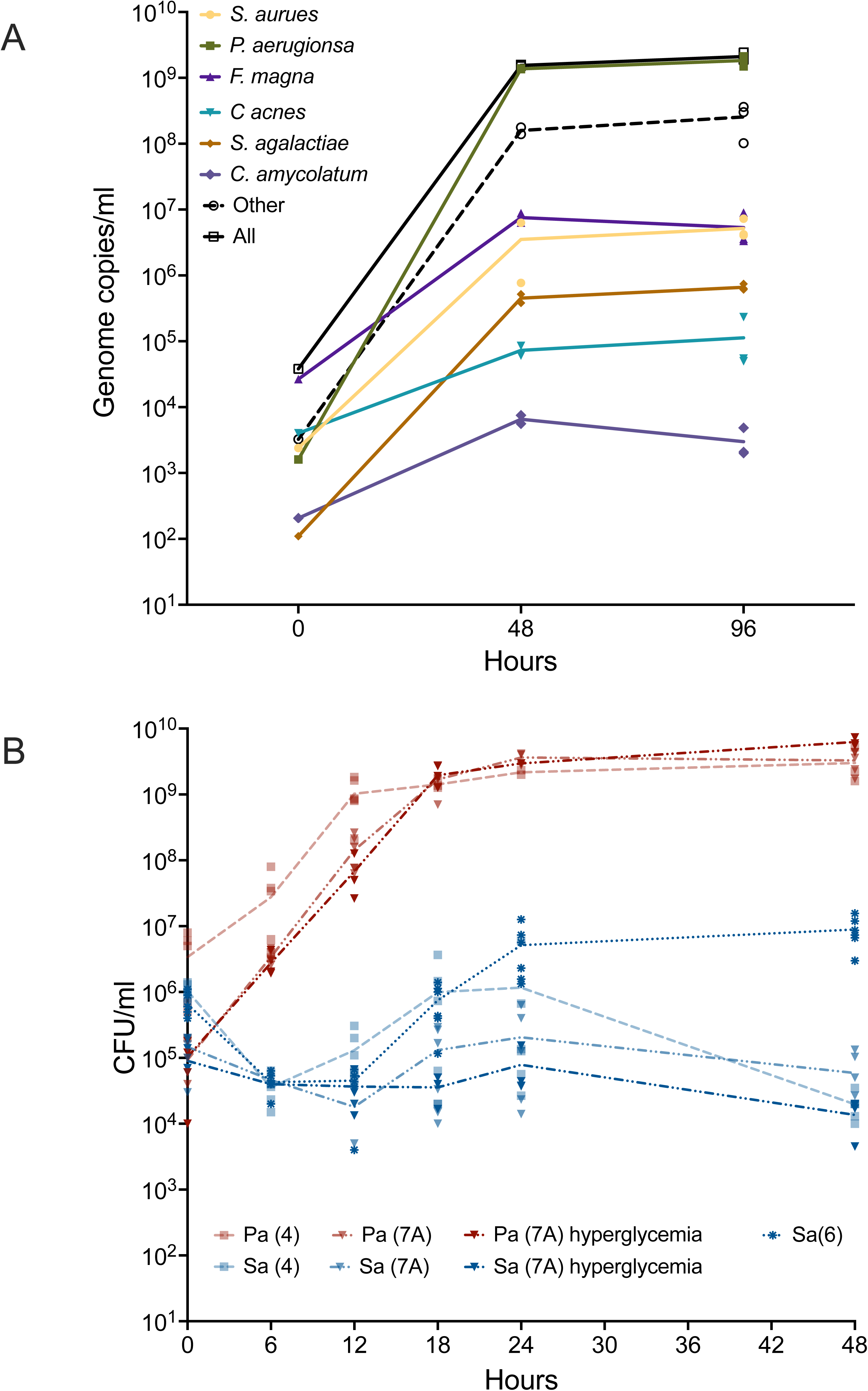

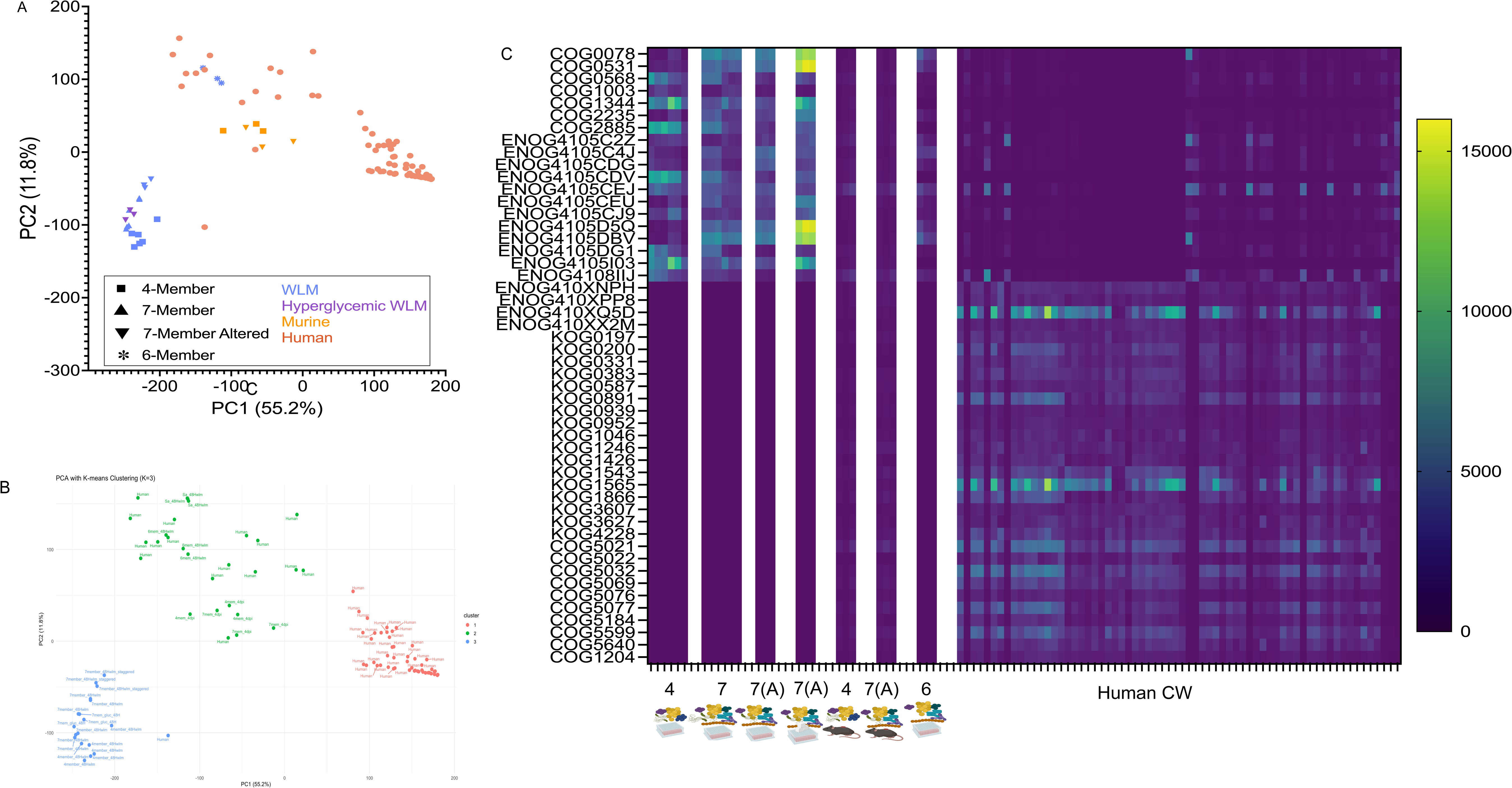

## Notes

### Competing Interest Statement

The authors have declared no competing interest.

