## Supplementary Information for "Revealing Community Dynamics in Polymicrobial Infections through a Quantitative Framework"

This PDF file includes:

Figures S1 to S4

Tables S1 to S4

Legends for Datasets S1 and S2

SI References

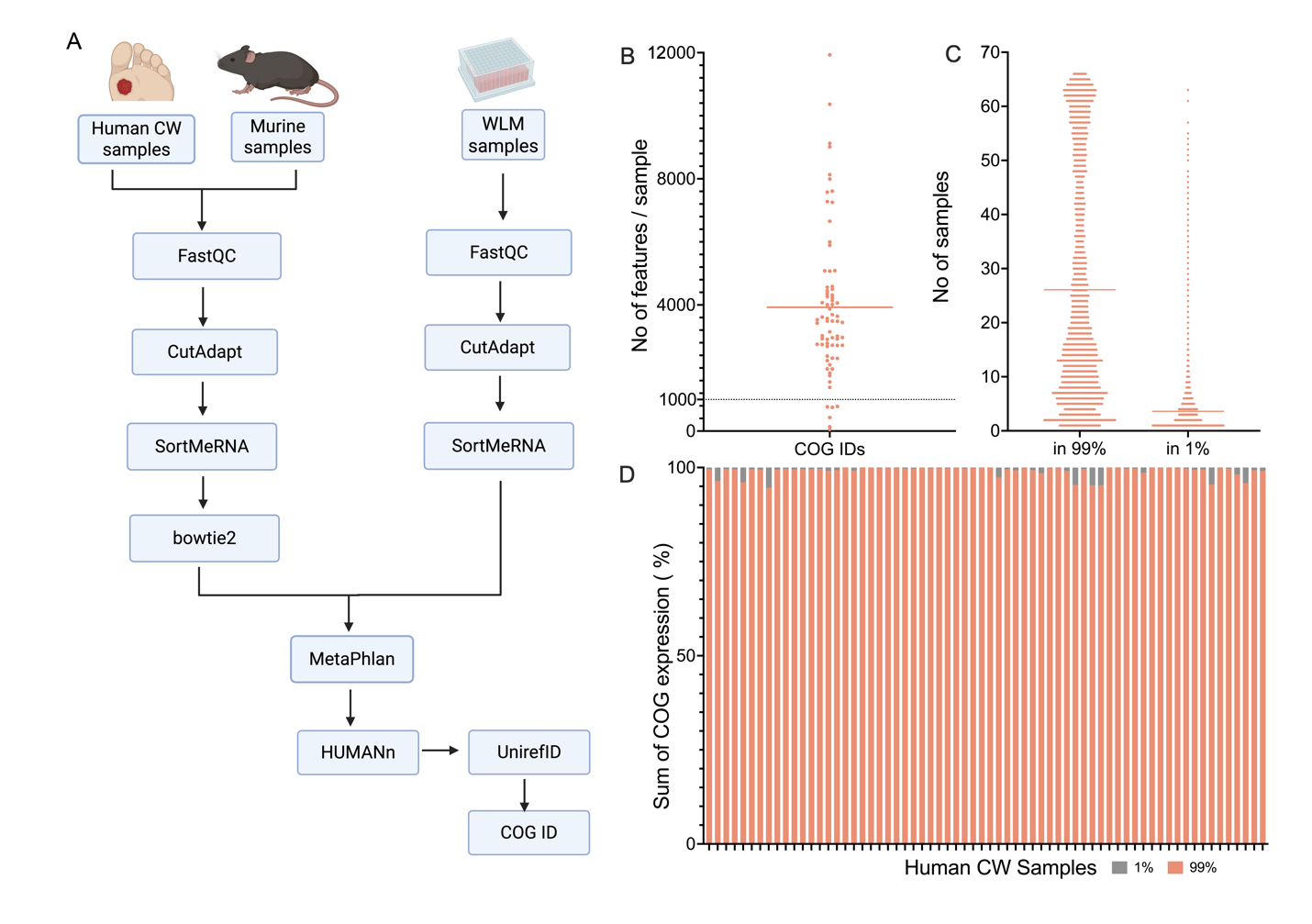

**Figure S1. Pipeline and quality control for inclusion of COG IDs in analysis pipeline. (A)** Graphical representation of bioinformatic steps for sample preprocessing and downstream analysis. **(B)** Distribution of COG IDs in human CW metatranscriptomic samples before inflection point analysis. Line at 1000 to highlight samples with less than 1000 COG IDs. **(C)** Representation of the number of samples that possess the COG IDs that make up the 99% (included in framework) and 1% (not included in framework) of expression data. Line at mean 26 and 4 for the 99% and 1% data, respectively. **(D)** How the COG IDs that make up the 99% and 1% of entire expression are distributed across the human CW samples.

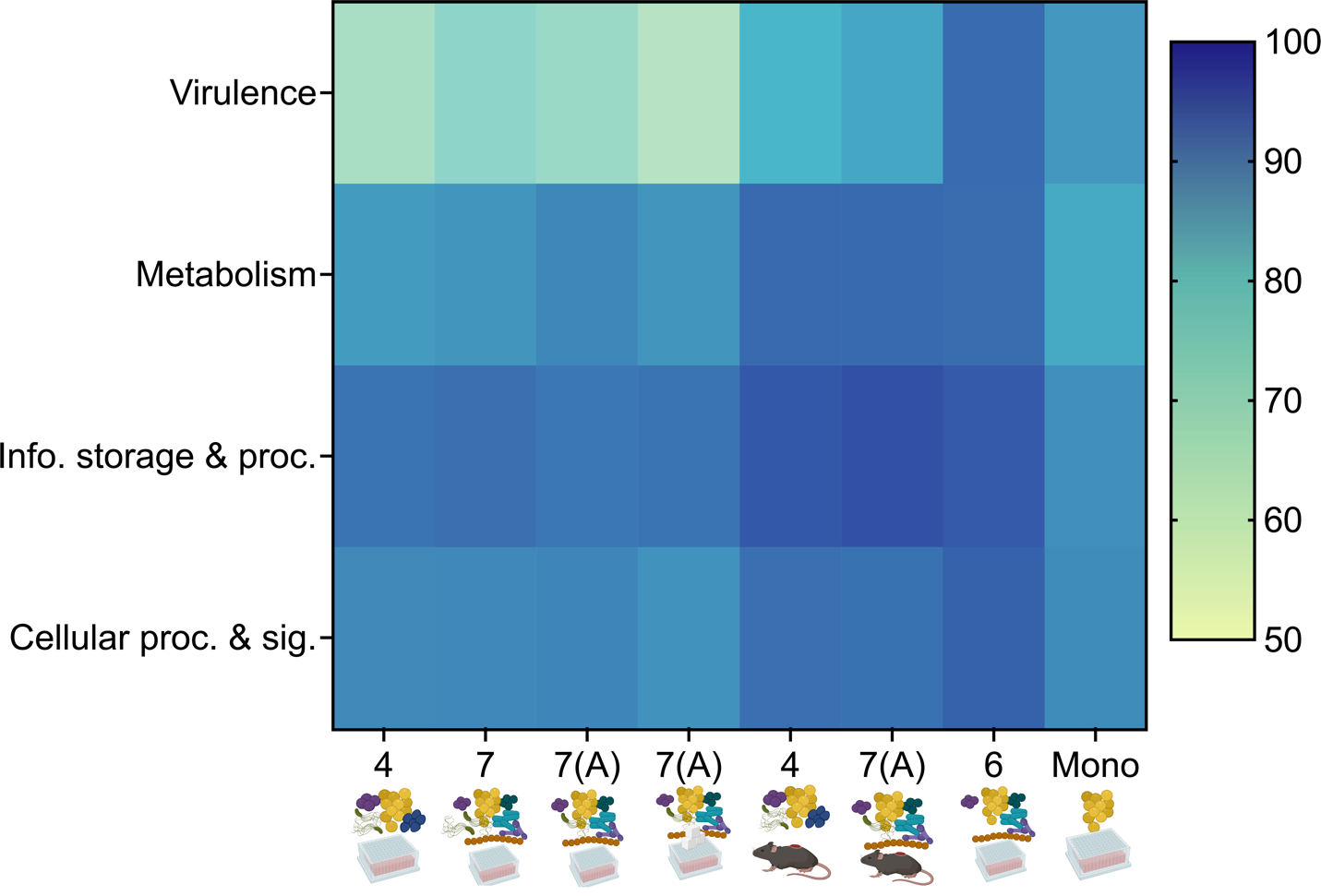

**Figure S2**: Virulence and cellar processes are highly impacted categories in the model communities. Graph shows distribution of the accuracy scores of the main categories in the samples evaluated with the overall accuracy scores being 4member_wlm 77.98%, 7member_wlm 78.00%, 7member_altered_wlm 82.02%, 7member_altered_hyperglycemia 77.62%, 4member_murine 89.70%, 6member_wlm 96.27%, *S. aureus* monoculture 89.80%. Abbreviations: 4 = 4member, 7 = 7member, 7(A) = 7member with altered inoculum, 6 = 6member, Mono = *S. aureus* monoculture in any condition.

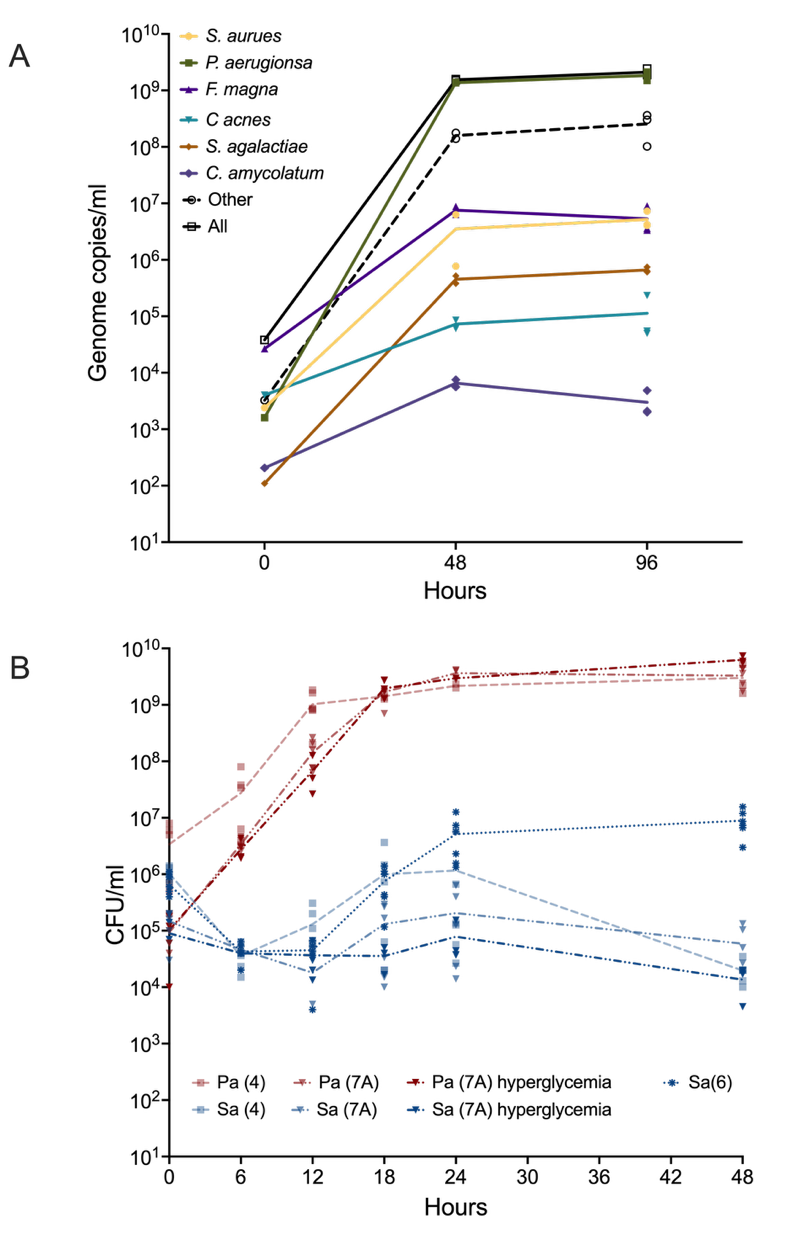

**Figure S3**: Growth of bacteria species in laboratory models**. (A)** qPCR results to show bacterial burden in the 7member community with altered inoculum in WLM. **(B)** Increase in the burden of *P. aeruginosa* shows a decrease in the burden of *S. aureus* in co-cultures. Graph shows the growth curve in WLM and hyperglycemic WLM to specifically track the growth of *S. aureus* (Sa) (blue) and *P. aeruginosa* (Pa) (red) in the four, six and seven member communities.

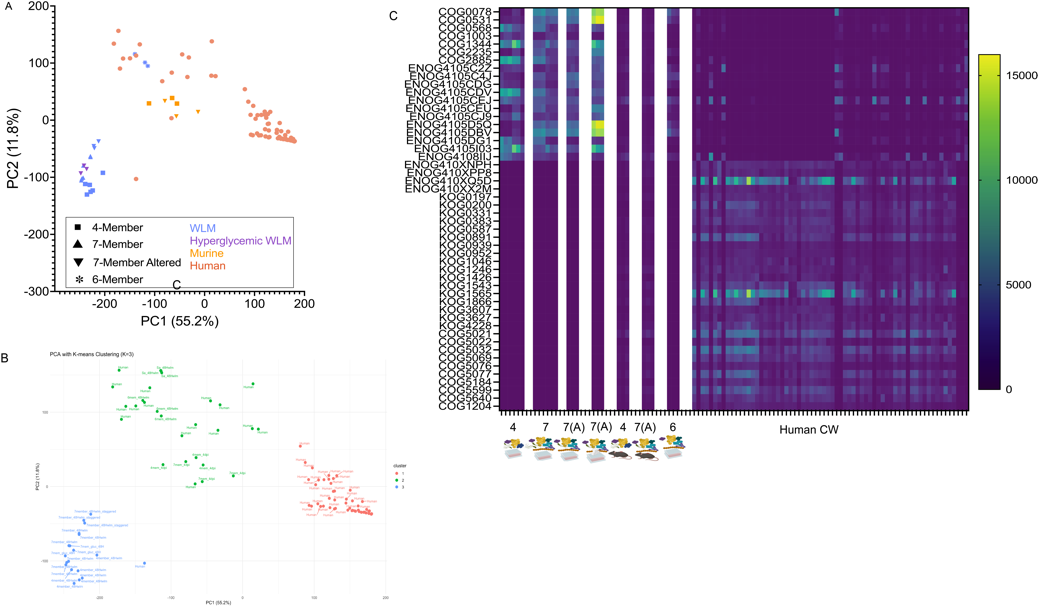

**Figure S4:** Six-member community (in vitro) and the in vivo four- and seven- member communities cluster more closely to some human samples. (A). PCA to show the clustering of samples. Samples are grouped as humans or model. Model samples are shaped by community structure and colored by infection environment. **(B)** Highlighting the clustering similarity using k-means clustering (**C)** Top 50 COG IDs that drive the clustering are expressed in similar patterns across the human, murine and invitro six member samples. Abbreviations: 4 = 4member, 7 = 7member, 7(A) = 7member with altered inoculum, 6 = 6member, Mono = *S. aureus* monoculture in any condition.

**Table S1.** Strains used in this study.

| Strain | Description | Identifier | Source or Reference |
| --- | --- | --- | --- |
| LAC | *S. aureus* LAC* (AH1263) community associated methicillin resistant USA300 isolate | CI3 | (1) |
| PAO1 | *P. aeruginosa* strain PAO1 wildtype strain | CI91 | (2) |
| SY01 | *F. magna* strain SY01 (HM-293), isolated from the vagina of a patient with bacterial vaginosis in Urbana, Illinois. Obtained from BEI Resources as part of the Human Microbiome Project. | CI201 | BEI Resources |
| V583 | *E. faecalis* vancomycin resistant isolate V583 | CI164 | (3) |
| A909 | *S. agalactiae* strain A909 isolated in 1934 from a septic human neonate. | CI185 | (4) |
| SK137 | *Cutibacterium acnes* (formerly *Propionibacterium acnes*) isolate SK137 (HM-122), isolated from the normal skin of a 57-year-old man. Obtained from BEI Resources as part of the Human Microbiome Project. | CI232 | BEI Resources |
| MJR7738A | *A. hydrogenalis* MJR7738A | CI233 | BEI Resources |
| SK46 | *C. amycolatum* strain SK46 (HM- 109D), isolated from the normal skin of a 57-year-old man. Obtained from BEI Resources as part of the Human Microbiome Project. | CI199 | BEI Resources |

**Table S2.** Quantitative PCR primers used in this study. All sequences are listed 5’ to 3’.

| *Species* | Forward | Reverse | Source |
| --- | --- | --- | --- |
| *P. aeruginosa* | TAAGGACAGCCAGGACTACGAGAA | TGGTAGATGGACGGTTCCCAGAAA | (5) |
| *S. aureus* | ATTTGGTCCCAGTGGTGTGGGTAT | GCTGTGACAATTGCCGTTTGTCGT | (5) |
| *F. magna* | TACTAATGAGAGTGGCGAACGGGT | ATTAATCCCGGTTTCCCGAGGCTA | (5) |
| *E. faecalis* | ACCAAGCGGCGTCAAGTATCAAGA | GTGTGCGCAATCGCTCCAATTTCT | (5) |
| All bacteria (Universal 16S) | CCATGAAGTCGGAATCGCTAG | GCTTGACGGGCGGTGT | (5) |
| *C. acnes* | GACATGGATCGGGAGTGCTC | CATAACGTGCTGGCAACAGTG | (6) |
| *S. agalactiae* | CACTCAGCTTGGGATAGATCAG | GAAGAGTCAGGTTCGGTCATT | This study |
| *C. amycolatum* | GGTTGATGTTGCGCTTTACC | GGCAGACTCTCGGAAAGAAA | This study |

**Table S3.** Metatranscriptomic samples used in this analysis. Red text indicate samples that were omitted from subsequent analyses following quality control. DFU = Diabetic Foot Ulcer

| **SRR ID** | **Source (Country)** | **City/State** | **Patient Age** | **Patient Sex** | **Infection type** | **Accession Number** |
| --- | --- | --- | --- | --- | --- | --- |
| SRR10074438 | Australia | Liverpool |  |  | DFU | PRJNA563930 |
| SRR10074439 | Australia | Liverpool | >18 | M | DFU | PRJNA563930 |
| SRR10074440 | Australia | Liverpool | 61 | F | DFU | PRJNA563930 |
| SRR10074441 | Australia | Liverpool | 68 | M | DFU | PRJNA563930 |
| SRR10074442 | Australia | Liverpool | 62 | M | DFU | PRJNA563930 |
| SRR10074443 | Australia | Liverpool |  |  | DFU | PRJNA563930 |
| SRR10074444 | Australia | Liverpool | 77 | M | DFU | PRJNA563930 |
| SRR10074445 | Australia | Liverpool | 71 | M | DFU | PRJNA563930 |
| SRR10074446 | Australia | Liverpool |  |  | DFU | PRJNA563930 |
| SRR10074447 | Australia | Liverpool |  |  | DFU | PRJNA563930 |
| SRR10074448 | Australia | Liverpool | 72 | M | DFU | PRJNA563930 |
| SRR10074449 | Australia | Liverpool |  |  | DFU | PRJNA563930 |
| SRR10074450 | Australia | Liverpool | 51 | F | DFU | PRJNA563930 |
| SRR10074451 | Australia | Liverpool | 68 | M | DFU | PRJNA563930 |
| SRR10074452 | Australia | Liverpool | 51 | F | DFU | PRJNA563930 |
| SRR10074453 | Australia | Liverpool | 64 | M | DFU | PRJNA563930 |
| SRR14174610 | Australia | Sydney | 60 | M | DFU | PRJNA720438 |
| SRR14174611 | Australia | Sydney | 54 | M | DFU | PRJNA720438 |
| SRR14174612 | Australia | Sydney |  |  | DFU | PRJNA720438 |
| SRR14174613 | Australia | Sydney | 54 | M | DFU | PRJNA720438 |
| SRR14174614 | Australia | Sydney |  |  | DFU | PRJNA720438 |
| SRR14174615 | Australia | Sydney |  |  | DFU | PRJNA720438 |
| SRR14174616 | Australia | Sydney | 69 | M | DFU | PRJNA720438 |
| SRR14174617 | Australia | Sydney | 38 | M | DFU | PRJNA720438 |
| SRR14174618 | Australia | Sydney |  |  | DFU | PRJNA720438 |
| SRR14174619 | Australia | Sydney |  |  | DFU | PRJNA720438 |
| SRR14174620 | Australia | Sydney |  |  | DFU | PRJNA720438 |
| SRR14174621 | Australia | Sydney | 54 | M | DFU | PRJNA720438 |
| SRR14174622 | Australia | Sydney | 54 | M | DFU | PRJNA720438 |
| SRR14174623 | Australia | Sydney | 51 | M | DFU | PRJNA720438 |
| SRR14174624 | Australia | Sydney | 51 | M | DFU | PRJNA720438 |
| SRR14174625 | Australia | Sydney | 51 | M | DFU | PRJNA720438 |
| SRR14174626 | Australia | Sydney |  |  | DFU | PRJNA720438 |
| SRR14174627 | Australia | Sydney |  |  | DFU | PRJNA720438 |
| SRR14174628 | Australia | Sydney |  |  | DFU | PRJNA720438 |
| SRR14174629 | Australia | Sydney | 38 | M | DFU | PRJNA720438 |
| SRR14374244 | Australia | Liverpool | 62 | M | DFU | PRJNA726011 |
| SRR14374245 | Australia | Liverpool | 68 | F | DFU | PRJNA726011 |
| SRR14374246 | Australia | Liverpool | 57 | M | DFU | PRJNA726011 |
| SRR14374247 | Australia | Liverpool | 56 | M | DFU | PRJNA726011 |
| SRR14374248 | Australia | Liverpool | 49 | M | DFU | PRJNA726011 |
| SRR14374249 | Australia | Liverpool | 46 | M | DFU | PRJNA726011 |
| SRR14374257 | Australia | Liverpool | 70 | M | DFU | PRJNA726011 |
| SRR14374258 | Australia | Liverpool | 42 | M | DFU | PRJNA726011 |
| SRR14374259 | Australia | Liverpool | 37 | M | DFU | PRJNA726011 |
| SRR14374260 | Australia | Liverpool |  |  | Unknown | PRJNA726011 |
| SRR14374261 | Australia | Liverpool | 36 | F | DFU | PRJNA726011 |
| SRR14374262 | Australia | Liverpool | 34 | F | DFU | PRJNA726011 |
| SRR14374263 | Australia | Liverpool | 34 | F | DFU | PRJNA726011 |
| SRR14374264 | Australia | Liverpool | 60 | M | DFU | PRJNA726011 |
| SRR14374265 | Australia | Liverpool | 69 | M | DFU | PRJNA726011 |
| SRR14374266 | Australia | Liverpool | 64 | M | DFU | PRJNA726011 |
| SRR14374267 | Australia | Liverpool | 46 | F | DFU | PRJNA726011 |
| SRR14374268 | Australia | Liverpool | 45 | M | DFU | PRJNA726011 |
| SRR14374269 | Australia | Liverpool | 64 | M | DFU | PRJNA726011 |
| SRR14374270 | Australia | Liverpool | 68 | M | DFU | PRJNA726011 |
| SRR14374271 | Australia | Liverpool | 51 | M | DFU | PRJNA726011 |
| SRR14374272 | Australia | Liverpool | 59 | M | DFU | PRJNA726011 |
| SRR14374273 | Australia | Liverpool | 56 | M | DFU | PRJNA726011 |
| SRR14374274 | Australia | Liverpool | 71 | M | DFU | PRJNA726011 |
| SRR14374275 | Australia | Liverpool | 71 | M | DFU | PRJNA726011 |
| SRR14374276 | Australia | Liverpool | 55 | M | DFU | PRJNA726011 |
| SRR14374277 | Australia | Liverpool | 67 | M | DFU | PRJNA726011 |
| SRR14374278 | Australia | Liverpool | 67 | M | DFU | PRJNA726011 |
| SRR14374279 | Australia | Liverpool | 64 | M | DFU | PRJNA726011 |
| SRR14374280 | Australia | Liverpool | 39 | M | DFU | PRJNA726011 |
| SRR6833323 | USA | Lubbock, TX | 36 | M | Foot Ulcer | SRP135669 |
| SRR6833324 | USA | Lubbock, TX | 88 | M | Foot Ulcer | SRP135669 |
| SRR6833325 | USA | Lubbock, TX | 31 | M | Foot Ulcer | SRP135669 |
| SRR6833340 | USA | Lubbock, TX | 59 | F | Foot Ulcer | SRP135669 |
| SRR6833343 | Denmark | Copenhagen | 69 | M | Foot Ulcer | SRP135669 |
| SRR6833348 | Denmark | Copenhagen |  |  | Foot Ulcer | SRP135669 |

**Table S4.** Metagenomic samples used in this analysis. DFU = Diabetic Foot Ulcer

| **SRRID** | **Source (Country)** | **City/State** | **Infection type** | **Accession Number** | **Diabetes Type** | **Age** | **Sex** |
| --- | --- | --- | --- | --- | --- | --- | --- |
| SRR8247667 | United States | Pennylvania | DFU | PRJNA506988 | Type 2 | 56 | Female |
| SRR8247668 | United States | Pennylvania | DFU | PRJNA506988 | Type 2 | 56 | Female |
| SRR8247671 | United States | Pennylvania | DFU | PRJNA506988 | Type 2 | 52 | Female |
| SRR8247672 | United States | Pennylvania | DFU | PRJNA506988 | Type 2 | 52 | Female |
| SRR8247673 | United States | Pennylvania | DFU | PRJNA506988 | Type 2 | 52 | Female |
| SRR8247674 | United States | Pennylvania | DFU | PRJNA506988 | Type 2 | 52 | Female |
| SRR8247675 | United States | Pennylvania | DFU | PRJNA506988 | Type 2 | 52 | Female |
| SRR8247676 | United States | Pennylvania | DFU | PRJNA506988 | Type 2 | 52 | Female |
| SRR8247677 | United States | Pennylvania | DFU | PRJNA506988 | Type 2 | 52 | Female |
| SRR8247678 | United States | Pennylvania | DFU | PRJNA506988 | Type 2 | 52 | Female |
| SRR8247679 | United States | Pennylvania | DFU | PRJNA506988 | Type 2 | 52 | Female |
| SRR8247682 | United States | Pennylvania | DFU | PRJNA506988 | Type 2 | 56 | Female |
| SRR8247683 | United States | Pennylvania | DFU | PRJNA506988 | Type 2 | 56 | Female |
| SRR8247686 | United States | Pennylvania | DFU | PRJNA506988 | Type 2 | 60 | Male |
| SRR8247689 | United States | Pennylvania | DFU | PRJNA506988 | Type 2 | 66 | Male |
| SRR8247690 | United States | Pennylvania | DFU | PRJNA506988 | Type 2 | 66 | Male |
| SRR8247693 | United States | Pennylvania | DFU | PRJNA506988 | Type 2 | 58 | Male |
| SRR8247694 | United States | Pennylvania | DFU | PRJNA506988 | Type 2 | 58 | Male |
| SRR8247695 | United States | Pennylvania | DFU | PRJNA506988 | Type 2 | 58 | Male |
| SRR8247696 | United States | Pennylvania | DFU | PRJNA506988 | Type 2 | 58 | Male |
| SRR8247697 | United States | Pennylvania | DFU | PRJNA506988 | Type 2 | 58 | Male |
| SRR8247698 | United States | Pennylvania | DFU | PRJNA506988 | Type 2 | 58 | Male |
| SRR8247699 | United States | Pennylvania | DFU | PRJNA506988 | Type 2 | 58 | Male |
| SRR8247700 | United States | Pennylvania | DFU | PRJNA506988 | Type 2 | 58 | Male |
| SRR8247701 | United States | Pennylvania | DFU | PRJNA506988 | Type 2 | 58 | Male |
| SRR8247704 | United States | Pennylvania | DFU | PRJNA506988 | Type 1 | 37 | Male |
| SRR8247707 | United States | Pennylvania | DFU | PRJNA506988 | Type 2 | 52 | Male |
| SRR8247708 | United States | Pennylvania | DFU | PRJNA506988 | Type 2 | 52 | Male |
| SRR8247711 | United States | Pennylvania | DFU | PRJNA506988 | Type 2 | 52 | Female |
| SRR8247712 | United States | Pennylvania | DFU | PRJNA506988 | Type 2 | 52 | Female |
| SRR8247713 | United States | Pennylvania | DFU | PRJNA506988 | Type 2 | 52 | Female |
| SRR8247714 | United States | Pennylvania | DFU | PRJNA506988 | Type 2 | 52 | Female |
| SRR8247715 | United States | Pennylvania | DFU | PRJNA506988 | Type 2 | 52 | Female |
| SRR8247716 | United States | Pennylvania | DFU | PRJNA506988 | Type 2 | 52 | Female |
| SRR8247717 | United States | Pennylvania | DFU | PRJNA506988 | Type 2 | 52 | Female |
| SRR8247718 | United States | Pennylvania | DFU | PRJNA506988 | Type 2 | 52 | Female |
| SRR8247721 | United States | Pennylvania | DFU | PRJNA506988 | Type 2 | 58 | Female |
| SRR8247722 | United States | Pennylvania | DFU | PRJNA506988 | Type 2 | 58 | Female |
| SRR8247725 | United States | Pennylvania | DFU | PRJNA506988 | Type 2 | 58 | Male |
| SRR8247728 | United States | Pennylvania | DFU | PRJNA506988 | Type 2 | 48 | Male |
| SRR8247729 | United States | Pennylvania | DFU | PRJNA506988 | Type 2 | 48 | Male |
| SRR8247732 | United States | Pennylvania | DFU | PRJNA506988 | Type 2 | 53 | Male |
| SRR8247733 | United States | Pennylvania | DFU | PRJNA506988 | Type 2 | 53 | Male |
| SRR8247736 | United States | Pennylvania | DFU | PRJNA506988 | Type 2 | 49 | Female |
| SRR8247737 | United States | Pennylvania | DFU | PRJNA506988 | Type 2 | 49 | Female |
| SRR8247738 | United States | Pennylvania | DFU | PRJNA506988 | Type 2 | 49 | Female |
| SRR8247739 | United States | Pennylvania | DFU | PRJNA506988 | Type 2 | 49 | Female |
| SRR8247740 | United States | Pennylvania | DFU | PRJNA506988 | Type 2 | 49 | Female |
| SRR8247741 | United States | Pennylvania | DFU | PRJNA506988 | Type 2 | 49 | Female |
| SRR8247742 | United States | Pennylvania | DFU | PRJNA506988 | Type 2 | 49 | Female |
| SRR8247743 | United States | Pennylvania | DFU | PRJNA506988 | Type 2 | 49 | Female |
| SRR8247748 | United States | Pennylvania | DFU | PRJNA506988 | Type 2 | 55 | Male |
| SRR8247751 | United States | Pennylvania | DFU | PRJNA506988 | Type 2 | 62 | Male |
| SRR8247754 | United States | Pennylvania | DFU | PRJNA506988 | Type 2 | 69 | Male |
| SRR8247759 | United States | Pennylvania | DFU | PRJNA506988 | Type 2 | 47 | Male |
| SRR8247764 | United States | Pennylvania | DFU | PRJNA506988 | Type 2 | 58 | Male |
| SRR8247765 | United States | Pennylvania | DFU | PRJNA506988 | Type 2 | 58 | Male |
| SRR8247766 | United States | Pennylvania | DFU | PRJNA506988 | Type 2 | 58 | Male |
| SRR11248549 | Australia | Liverpool | DFU | PRJNA610303 | Type 1 | 54 | Male |
| SRR11248550 | Australia | Liverpool | DFU | PRJNA610303 | Type 1 | 51 | Female |
| SRR11248551 | Australia | Liverpool | DFU | PRJNA610303 | Type 1 | 59 | Male |
| SRR11248552 | Australia | Liverpool | DFU | PRJNA610303 | Type 2 | 54 | Male |
| SRR11248553 | Australia | Liverpool | DFU | PRJNA610303 | Type 2 | 63 | Male |
| SRR11248554 | Australia | Liverpool | DFU | PRJNA610303 | Type 2 | 50 | Female |
| SRR11248555 | Australia | Liverpool | DFU | PRJNA610303 | Type 2 | 65 | Male |
| SRR11248556 | Australia | Liverpool | DFU | PRJNA610303 | Type 2 | 88 | Female |
| SRR11248557 | Australia | Liverpool | DFU | PRJNA610303 | Type 2 | 77 | Male |
| SRR11248558 | Australia | Liverpool | DFU | PRJNA610303 | Type 2 | 50 | Male |
| SRR11248559 | Australia | Liverpool | DFU | PRJNA610303 | Type 2 | 82 | Female |
| SRR11248560 | Australia | Liverpool | DFU | PRJNA610303 | Type 2 | 62 | Male |
| SRR11248561 | Australia | Liverpool | DFU | PRJNA610303 | Type 2 | 53 | Male |
| SRR11248562 | Australia | Liverpool | DFU | PRJNA610303 | Type 2 | 61 | Female |
| SRR11248563 | Australia | Liverpool | DFU | PRJNA610303 | Type 2 | 71 | Male |
| SRR11248564 | Australia | Liverpool | DFU | PRJNA610303 | Type 2 | 54 | Male |
| SRR11248565 | Australia | Liverpool | DFU | PRJNA610303 | Type 2 | 67 | Female |
| SRR11248566 | Australia | Liverpool | DFU | PRJNA610303 | Type 2 | 61 | Female |
| SRR11248567 | Australia | Liverpool | DFU | PRJNA610303 | Type 1 | 51 | Female |
| SRR11248568 | Australia | Liverpool | DFU | PRJNA610303 | Type 1 | 57 | Male |
| SRR11248569 | Australia | Liverpool | DFU | PRJNA610303 | Type 2 | 51 | Male |
| SRR11248570 | Australia | Liverpool | DFU | PRJNA610303 | Type 2 | 64 | Male |
| SRR11248571 | Australia | Liverpool | DFU | PRJNA610303 | Type 2 | 67 | Male |
| SRR11248572 | Australia | Liverpool | DFU | PRJNA610303 | Type 2 | 72 | Male |
| SRR11248573 | Australia | Liverpool | DFU | PRJNA610303 | Type 2 | 71 | Male |
| SRR11248574 | Australia | Liverpool | DFU | PRJNA610303 | Type 2 | 54 | Male |
| SRR11248575 | Australia | Liverpool | DFU | PRJNA610303 | Type 2 | 67 | Male |
| SRR11248576 | Australia | Liverpool | DFU | PRJNA610303 | Type 2 | 69 | Male |
| SRR11248577 | Australia | Liverpool | DFU | PRJNA610303 | Type 2 | 68 | Male |
| SRR11248578 | Australia | Liverpool | DFU | PRJNA610303 | Type 2 | 54 | Male |
| SRR11248579 | Australia | Liverpool | DFU | PRJNA610303 | Type 2 | 81 | Female |
| SRR11248580 | Australia | Liverpool | DFU | PRJNA610303 | Type 2 | 66 | Male |
| SRR11248581 | Australia | Liverpool | DFU | PRJNA610303 | Type 1 | 48 | Male |
| SRR11248582 | Australia | Liverpool | DFU | PRJNA610303 | Type 1 | 36 | Female |
| SRR11248583 | Australia | Liverpool | DFU | PRJNA610303 | Type 2 | 68 | Male |
| SRR11248584 | Australia | Liverpool | DFU | PRJNA610303 | Type 2 | 64 | Male |

**Dataset S1 (separate file):** Distribution of functions for each annotation scheme in each sample. **A**: Column A to G showing all the human samples and their associated number of features from different annotations; Column J showing the 6 human samples that were removed prior to inflection point analysis. **B**: Showing emphasis on samples whose UniRef IDs were lesser than GO terms when collapsed. **C**: Showing the inflection point analysis result and how many IDs make 99%. **D**: Showing the distribution of the COG IDs after inflection point analysis. **E**: Showing the distribution of COG IDs in all the samples with their mean.

**Dataset 2 (separate file):** Taxonomic data from MetaPhlAn4 for metatranscriptomic and metagenomic datasets. Cumulative abundance data of species in the metatranscriptomic (**A**) and metagenomic (**B**) datasets also showing the top 80% and overlaps between the two datasets (**C**).
